# SHADE: A Multilevel Bayesian Approach to Modeling Directional Spatial Associations in Tissues

**DOI:** 10.1101/2025.06.24.661393

**Authors:** Joel Eliason, Michele Peruzzi, Arvind Rao

## Abstract

**Motivation:** Spatial dependencies in tissue microenvironments, particularly asymmetric interactions between cell types, are central to understanding immune dynamics, tumor behavior, and tissue organization. Existing spatial statistical methods often assume symmetric associations or analyze images independently, limiting biological interpretability and inference quality.

**Results:** We introduce SHADE (Spatial Hierarchical Asymmetry via Directional Estimation), a Bayesian hierarchical framework that models asymmetric spatial associations and multilevel structure in multiplexed imaging data. SHADE captures directional relationships via smooth spatial interaction curves (SICs), provides interpretable distance-resolved summaries of cell-cell interactions, and supports multiscale inference across tissue sections, patients, and cohorts. Simulation studies demonstrate improved inference quality and robustness, and application to colorectal cancer imaging data reveals biologically meaningful differences in immune and stromal organization.

**Availability and Implementation:** Source code and analysis scripts are freely available at http://github.com/jeliason/SHADE and http://github.com/jeliason/shade_paper_code, implemented in R and Stan.

## 1 Introduction

Spatial dependencies between cell types play a central role in immune dynamics, tumor behavior, and tissue organization, motivating statistical models that can capture such interactions [Yuan, 2016, Maley et al., 2017]. The tumor microenvironment (TME) is a spatially structured system where cell arrangements are closely linked to disease progression and treatment response. Notably, these spatial interactions are frequently asymmetric: the spatial pattern of one cell type may be associated with that of another without necessarily implying reciprocity. For example, anti-tumor immune cells often cluster near tumor cells as part of immune surveillance, while tumor positioning is largely determined by vasculature and tissue architecture rather than by the presence of immune cells [Binnewies et al., 2018, Gilkes et al., 2014, Bindea et al., 2013]. Quantifying such asymmetric associations is important for understanding tissue-level biological processes.

Recent advances in multiplexed imaging technologies, including multiplexed immunofluorescence (mIF) and other high-resolution spatial profiling methods, have made it possible to quantify cell type spatial distributions and interactions within the tumor microenvironment (TME) at single-cell resolution [Sheng et al., 2023]. However, conventional spatial statistics, such as Ripley’s *K*-function and pair correlation functions, primarily describe co-location patterns and do not explicitly model conditional relationships. Although Gibbs point process models offer a more flexible probabilistic framework [Moller and Waagepetersen, 2003, Baddeley et al., 2015], standard formulations assume symmetric interactions, thereby overlooking the possibility that one cell type may be associated with a directional spatial arrangement of another. While hierarchical Gibbs models have been proposed to accommodate directionality [Högmander and Särkkä, 1999, Grabarnik and Särkkä, 2009], these methods often depend on parametric interaction functions that impose restrictive assumptions.

Observations based on the *G*-cross function suggest that spatial interactions between cell types are inherently asymmetric, with the inferred dependence structure varying by the choice of reference [Tsang et al., 2024]. These findings motivate statistical models that directly account for directional spatial effects specified by biological hypotheses.

In this work, we develop a flexible statistical framework for modeling asymmetric spatial associations in tissue microenvironments, making it well-suited for the analysis of multiplexed imaging datasets. While our approach is not a direct extension of the multitype Gibbs point process (MGPP) model (see Eliason and Rao [2024], Flint et al. [2022]), it is certainly inspired by it and was developed specifically to address some of its limitations—particularly the inability to handle asymmetric interactions and to incorporate multilevel structure. Furthermore, our framework extends the aforementioned hierarchical Gibbs models by incorporating *spatial interaction curves (SICs)*, which flexibly quantify how the presence of one cell type relates to the expected density of another across different spatial scales. This is achieved by modeling the *conditional intensity function*, which describes the expected density of a target cell type at any spatial location given the observed pattern of a reference cell type. Unlike traditional step-function interaction terms, we model SICs via a flexible spline basis, enabling smooth, data-driven representations of spatial associations without imposing arbitrary interaction radii or functional forms—while maintaining interpretability through an explicit model of how local cell configurations contribute to overall spatial organization.

Beyond asymmetry, our method accounts for multilevel variation across images, patients, and cohorts. Biological spatial patterns vary at multiple levels, yet existing spatial models typically analyze images independently, potentially overlooking important heterogeneity across patients. We integrate the estimation of SICs into a multilevel Bayesian framework, which enables partial pooling across biological scales while preserving biologically meaningful variation.

To ensure computational efficiency, we approximate conditional intensity estimation using logistic regression with quadrature-based dummy points rather than direct Poisson likelihood estimation [Baddeley et al., 2014]. This transformation avoids computational challenges such as instability in fine-grained quadrature approximations and biases inherent in conventional spatial regression techniques.

We refer to this overall modeling framework as SHADE (Spatial Hierarchical Asymmetry via Directional Estimation). SHADE combines asymmetric spatial association modeling with hierarchical Bayesian inference to capture complex spatial organization across biological scales. An overview of the SHADE workflow is provided in Figure 1.

**Figure 1:**
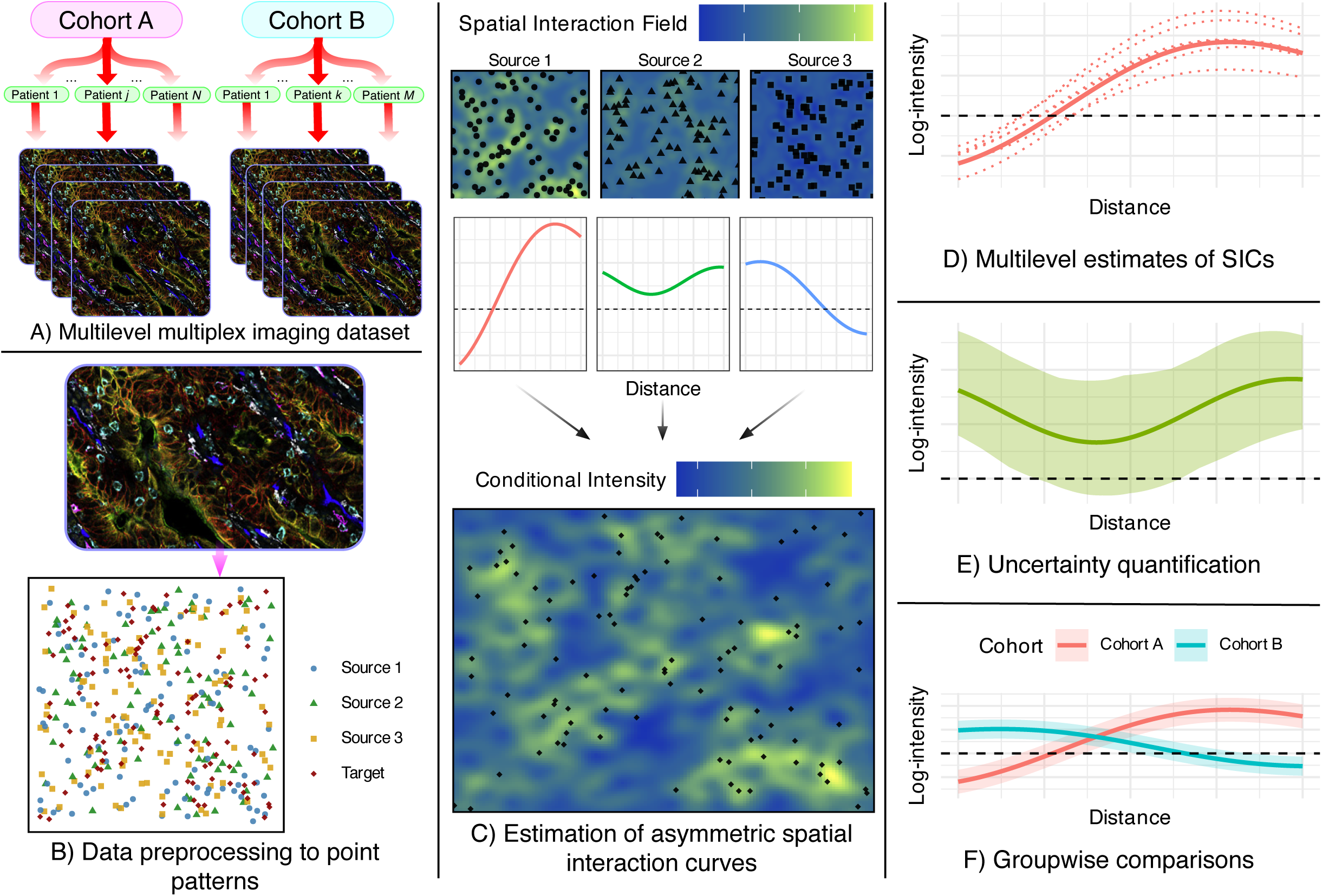
Summary of the SHADE (Spatial Hierarchical Asymmetry via Directional Estimation) framework. **A)** Multiplexed imaging data is structured hierarchically across cohorts, patients, and images. Multiplex images were adapted from Schürch et al. [2020] and are used under a Creative Commons CC BY 4.0 license. **B)** Images are processed into spatial point patterns with cell type annotations. **C)** SHADE estimates Spatial Interaction Curves (SICs) that capture directional associations between cell types across spatial scales. **D)** SICs are estimated at cohort, patient, and image levels, enabling multilevel analysis of spatial heterogeneity. **E)** Posterior distributions provide uncertainty quantification. **F)** SICs can be compared across cohorts to assess differences in spatial organization.

Our approach builds on existing spatial point process models while addressing several key limitations in the analysis of tissue microenvironments. First, we introduce a conditional intensity modeling framework that explicitly captures directional spatial associations between cell types, enabling the study of asymmetric interactions. Second, we develop a nonparametric representation of spatial interactions through spatial interaction curves, which provide a flexible and interpretable summary of how cell types influence one another as a function of distance. Third, we incorporate a multilevel Bayesian structure that models variation in spatial interactions across images, patients, and cohorts, allowing for partial pooling and uncertainty quantification at each level. Finally, we adopt a logistic regression approximation to the Poisson likelihood, which enables scalable and numerically stable inference without the need for fine spatial discretization.

## 2 Methods

### 2.1 Hierarchical Modeling of Conditional Spatial Point Processes

We model the spatial distribution of a target cell type *B* given the presence of one or more conditioning cell types *A*_1_*, A*_2_*, . . . , A_K_*, using a hierarchical framework based on conditional spatial point processes. Our approach extends hierarchical Gibbs models [Högmander and Särkkä, 1999, Grabarnik and Särkkä, 2009] and is designed to flexibly estimate asymmetric spatial association patterns while accounting for multilevel variation across images, patients, and cohorts.

Let *X_A_k__, X_B_* ⊂ *W* denote the observed spatial point patterns of cell types *A_k_* and *B*, respectively, within a two-dimensional tissue region *W* ⊂ ℝ^2^. Formally, 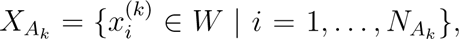 *X_B_* = {*y_j_* ∈ *W* | *j* = 1*, . . . , N_B_*}, where 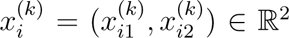 denotes the two-dimensional spatial coordinate of the *i*-th cell of type *A_k_*, and *y_j_* = (*y_j_*_1_*, y_j_*_2_) ∈ ℝ^2^ denotes the coordinate of the *j*-th cell of type *B*. The quantities *N_A_k__* and *N_B_* indicate the total number of observed cells of type *A_k_* and *B*, respectively. The observation window *W* corresponds to the region of tissue captured in the image, typically a rectangular subset of the plane defined by the image dimensions.

In practice, *X_A_k__* and *X_B_* are obtained from image segmentation and cell type classification pipelines applied to high-resolution tissue images, such as those generated by multiplexed imaging platforms.

We model the spatial distribution of *X_B_* (the *target* cell type, e.g., immune cells) conditional on *X*_*A*_1__ *, . . . , X_A_K__* (the *source* cell types, e.g., tumor cells and vasculature) by assuming that *X_B_* | *X*_*A*_1__ *, . . . , X_A_K__* follows an inhomogeneous Poisson point process [Baddeley et al., 2015], the likelihood of which we write as:

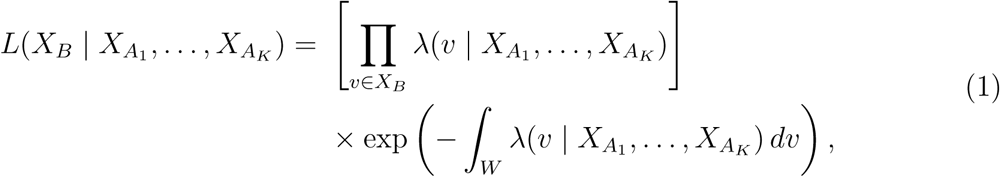

where the product runs over all observed locations of type *B*, while the integral accounts for the total expected intensity over the observation window *W* . The point process for *X_B_* is entirely characterized by its *conditional intensity function λ*(*v* | *X_A_*_1_ *, . . . , X_A_K__*), which defines the expected local density of type *B* cells at any location *v* ∈ *W*, conditioned on the spatial configurations of the conditioning cell types, that is, 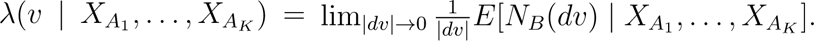

We model *λ*(·) as depending log-linearly on the observed patterns of *A*_1_*, . . . , A_K_* cells:

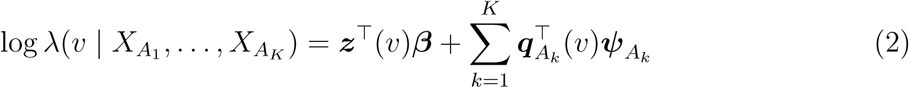

In (2), the term ***z***(*v*) ∈ ℝ*^J^* is a vector of covariates at location *v* (including an intercept), with corresponding coefficients ***β*** ∈ ℝ*^J^* . The vector ***q****_A_* (*v*) ∈ ℝ*^P^* encodes spatial interaction features between type *A_k_* and type *B*; its *p*-th element is defined as:

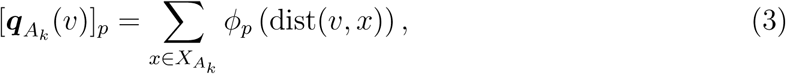

where *ϕ_p_*(·) is a basis function (e.g., a B-spline or Gaussian kernel) that modulates the influence of a type *A_k_* cell at a given distance from *v*. Finally, ***ψ****_A_k__* ∈ ℝ*^P^* are the corresponding coefficients. This formulation flexibly captures how proximity to different cell types influences the expected density of type *B* cells across spatial scales. Our use of smooth interaction features in (3) reflects biologically realistic, distance-dependent associations. In doing so, we generalize traditional Gibbs process formulations by avoiding rigid parametric forms such as fixed interaction radii [Grabarnik and Särkkä, 2009], which fail to capture the continuous and often subtle variations observed in biological spatial interactions [Baddeley and Turner, 2005].

While the model specified in (2) provides a flexible representation of spatial interactions through basis expansions, directly interpreting the estimated coefficients ***ψ****_A_k__* can be difficult. In particular, biological interest often centers on how the strength and direction of association between a conditioning cell type *A_k_* and a target cell type *B* vary as a function of distance. To facilitate interpretable biological insights, we next define a derived quantity, the *spatial interaction curve* (SIC), which summarizes the estimated effect of proximity to *A_k_* cells on the expected density of *B* cells across spatial scales.

#### 2.1.1 Spatial Interaction Curve

The SIC summarizes the asymmetric spatial association between a conditioning cell type *A_k_* and a target cell type *B* as a function of distance *s*:

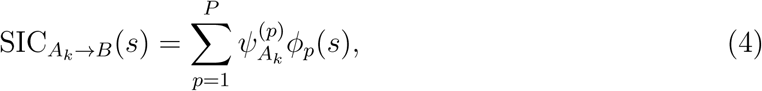

This curve represents the expected contribution of type *A_k_* cells to the log-intensity of type *B* cells as a function of distance *s* from a type *A_k_* cell.

As illustrated in Figure 2, a positive SIC value at distance *s* indicates that the presence of a type *A_k_* cell is associated with an increased expected density of type *B* cells at that radius, consistent with spatial attraction or clustering. Conversely, a negative SIC value suggests spatial repulsion or avoidance at that distance. This function provides an interpretable, distance-dependent summary of directional spatial association between cell types. Additional explanation of the SIC is contained in the Supplement.

**Figure 2:**
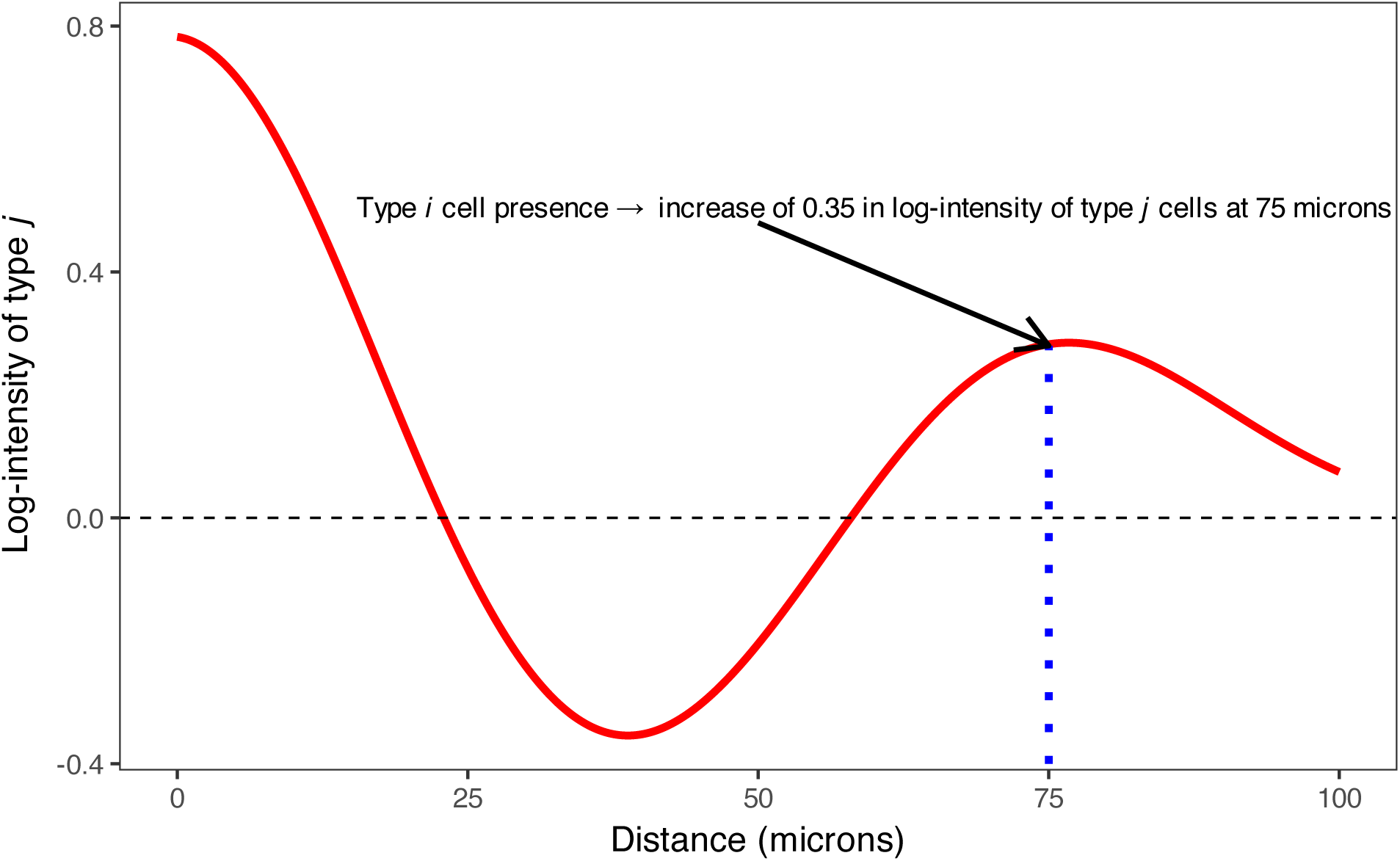
An example spatial interaction curve showing the effect of a source cell type *A_k_* on a target cell type *B*. At each distance *s*, the curve value represents the change in log-intensity of type *B* associated with the presence of a type *A_k_* cell at distance *s*.

A key advantage of our framework is that the spatial interaction terms ***ψ****_A_k__* can take both positive and negative values, allowing for flexible modeling of attraction and repulsion. This mirrors the flexibility of hierarchical Gibbs models while overcoming constraints in symmetric pairwise interaction models, which typically restrict interaction terms to be non-positive and therefore cannot directly model spatial attraction.

#### 2.1.2 Characterizing Predictive Asymmetry

Our modeling framework captures *predictive* spatial associations, not biological causation. A strong *A* → *B* SIC indicates that the presence of *A* is statistically predictive of local *B* density, but it does not imply that *A* exerts a causal biological effect on *B*. Apparent directional associations may arise from confounding factors such as differences in the spatial distribution of *A* (e.g., sparser, more clustered, or more localized patterns), which can make *A* a stronger statistical predictor without implying direct biological influence.

To mitigate such confounding, the model can incorporate spatial covariates or trend surfaces (e.g., distance to the tumor margin) to adjust for first-order intensity effects. This adjustment ensures that estimated directional SICs capture residual spatial association beyond shared background structure.

### 2.2 Multilevel Bayesian Model

To capture biological variability across individuals and sampling levels, we impose a hierarchical prior structure on the spatial interaction coefficients. For each conditioning cell type *A_k_*, the interaction coefficients are indexed hierarchically across three levels: cohort (*ψ*), patient (*γ*), and image (*δ*). Specifically, for each basis function *p*, we define:

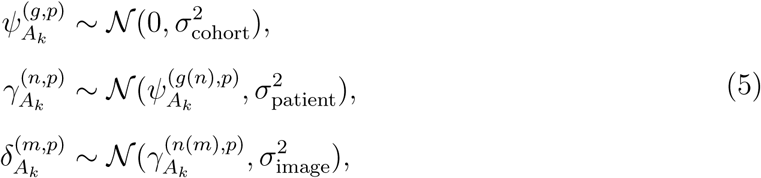

where *g*(*n*) maps patient *n* to its cohort, and *n*(*m*) maps image *m* to its corresponding patient. Each image-level coefficient 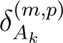 governs the localized effect of source cell type *A_k_* on target cell type *B* at distance scale *p*, within image *m*.

This hierarchical formulation enables partial pooling across the dataset structure, improving estimation stability while preserving biologically meaningful heterogeneity in spatial interactions. By modeling variation at the cohort, patient, and image levels, the framework supports inference on both shared and context-specific spatial association patterns.

Hyperpriors for the variance components 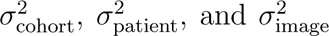 are detailed in the Supplement.

### 2.3 Logistic Regression Approximation for Computational Efficiency

Direct estimation of the Poisson likelihood in spatial point process models typically requires numerical integration over a fine spatial grid, which becomes computationally expensive and unstable in high-resolution images. To address this, we follow the logistic regression approximation introduced by Baddeley et al. [2014], which avoids spatial gridding by introducing a set of dummy points *D* sampled from a homogeneous Poisson process with known intensity *λ*_dummy_. We define the combined set of points *Y* = *X_B_* ∪ *D*, and assign binary labels:

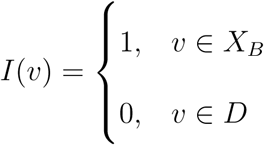

The conditional probability that a point *v* is a true (observed) point of type *B* is given by:

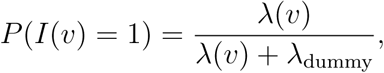

leading to the following logistic regression model:

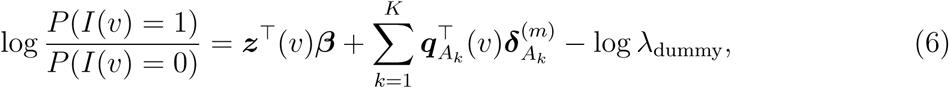

This approximation circumvents key computational challenges of direct Poisson modeling. In Poisson models, fine spatial discretization leads to large numbers of empty pixels, often resulting in singular design matrices and unstable inference [Baddeley et al., 2015]. The Hauck-Donner effect [Hauck and Donner, 1977] can further distort uncertainty estimates. By reframing the problem as a binary classification task over observed and dummy points, the logistic approximation enables scalable, stable inference of spatial interaction effects at the image level.

### 2.4 Summary of Estimation Procedure

Model fitting is implemented in `Stan` using Hamiltonian Monte Carlo (HMC) via `cmdstanr` [Stan Development Team, 2024, Gabry et al., 2024].

To approximate the likelihood, we first generate a set of dummy points *D* from a homogeneous Poisson process with intensity *λ*_dummy_ over the observation window *W* . For each location *v* ∈ *X_B_* ∪ *D*, we then compute spatial covariates ***z***(*v*) and interaction features ***q****_A_k__* (*v*), where each interaction feature encodes basis-function-weighted distances to cells of type *A_k_*.

Using these constructed features, we fit a multilevel Bayesian logistic regression model based on the approximation in (6), with spatial interaction coefficients 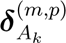 modeled hierarchically according to the priors in (5). Finally, we extract posterior draws of the interaction coefficients and reconstruct the spatial interaction curves using (4).

## 3 Simulation Studies

To evaluate the proposed model, we conducted simulation studies generating synthetic spatial point patterns that incorporated asymmetric interactions and multilevel structure. Spatial patterns were simulated within a bounded domain *W* = [0*, S*] × [0*, S*], with source points generated from a homogeneous Poisson process and target points influenced by spatial interaction curves defined over source-target distances. Hierarchical structure was introduced via interaction coefficients generated according to (5).

The final point pattern was converted into a logistic regression dataset using dummy points sampled from a homogeneous Poisson process with intensity *λ*_dummy_, and spatial interaction features were computed as in (3). Full simulation details, including two additional studies exploring the effect of various hyperparameters on statistical inference, are provided in the Supplement.

### 3.1 The effect of explicitly modeling multilevel structure

In our first simulation study, we examined how accounting for hierarchical structure affects inference quality. We simulated data from two patient groups, each with 20 patients and 4 images per patient. Each image contained 150 points of each type, with a dummy-to-real point ratio of 2. The remaining parameters were set to defaults (see Supplement).

We generated 100 simulation replicates and fit two models to each: the full hierarchical model and a non-hierarchical model that estimates image-level SICs independently, without shrinkage to patient- or cohort-level structures. Inference quality was assessed by comparing the average RMSE of the *δ*^(*m,p*)^ coefficients.

We found that on average, and across all spatial scales, the average RMSE of *δ*^(*m,p*)^ coefficients was lower for those estimated by the full hierarchical model than the model that fit images individually, most significantly so for closer-range coefficients (Table 1).

**Table 1:** Average RMSE of *δ*^(*m,p*)^ coefficients in aggregate and across spatial scales, from both a model that fully accounts for the hierarchical structure as well as a model that does not account for hierarchy.

| Hierarchical | Scale: All | Scale: Small | Scale: Medium | Scale: Large |
| --- | --- | --- | --- | --- |
| False | 0.142 (0.097, 0.196) | 0.355 (0.230, 0.509) | 0.022 (0.016, 0.031) | 0.048 (0.034, 0.066) |
| True | 0.039 (0.027, 0.043) | 0.072 (0.043, 0.084) | 0.017 (0.013, 0.020) | 0.028 (0.021, 0.036) |

Figure 3 shows several examples of image-level SICs estimated with both models from this simulation study. In all cases, we can see that the SIC is better estimated when hierarchical structure is accounted for, in terms of both the bias and and variance of the estimate.

**Figure 3:**
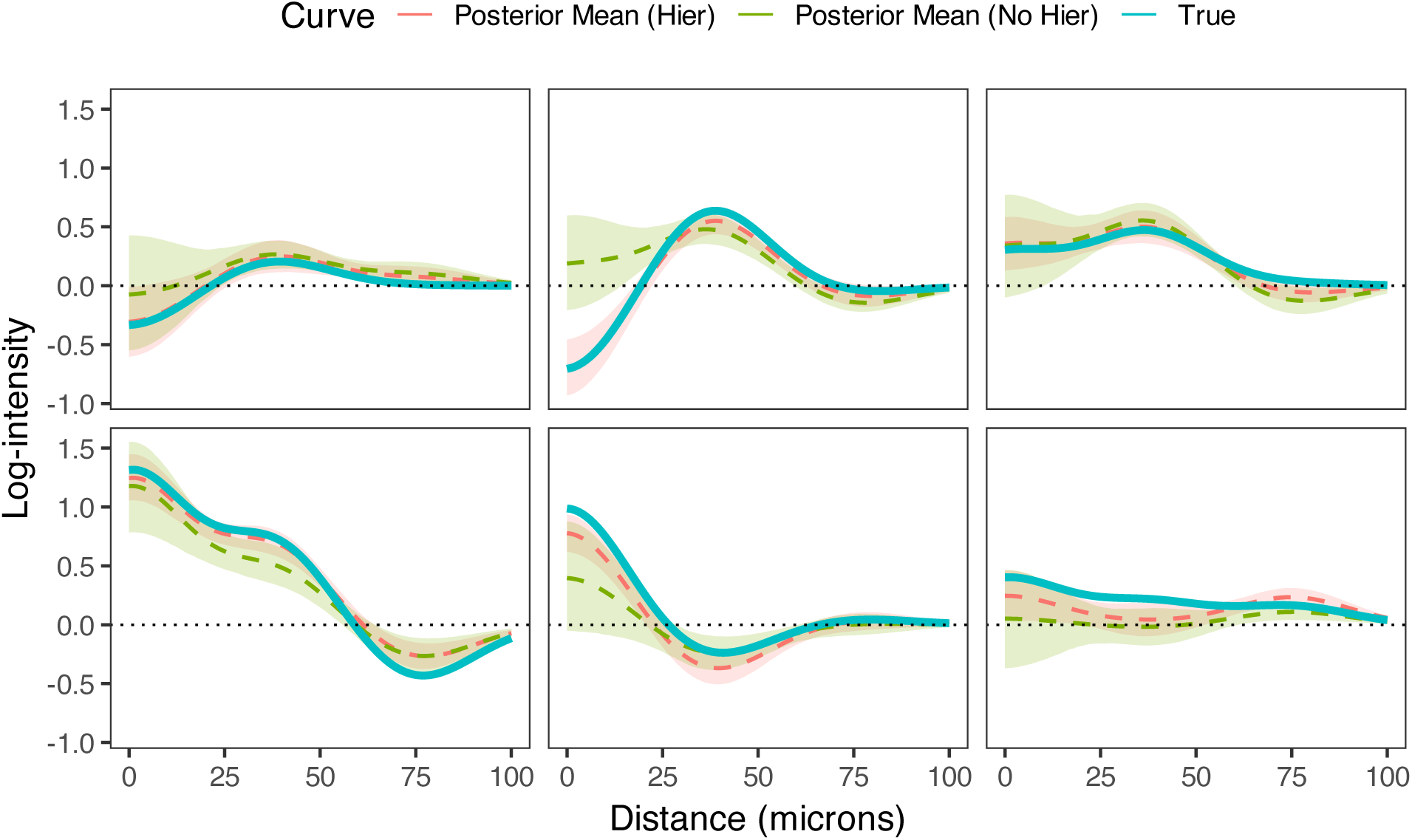
Examples of estimated image-level SICs from hierachical modeling simulations, demonstrating better estimation of SICs when accounting for multilevel structure.

### 3.2 Comparison of spatial pattern detection accuracy across methods

We also conducted a simulation study to evaluate SHADE’s performance against established spatial analysis methods. We generated hierarchical spatial point patterns with known asymmetric interactions between T cells and tumor cells, simulating responder and non-responder patient groups with distinct spatial signatures. Responder patterns exhibited strong spatial clustering (positive spatial interaction curves), while non-responder patterns showed strong repulsion (negative curves).

We simulated spatial point patterns with known group-, patient-, and image-level structure, using radial basis functions to model short-, medium-, and long-range interactions. T cell and tumor cell densities (15 vs. 150) and the number of images per patient (1–3) were varied across 30 replicates per condition. We evaluated SHADE’s ability to recover the sign of spatial interactions at target distances (20, 40, 60 *µ*m) using pointwise 95% credible intervals. For comparison, we applied *G*-cross envelope tests based on 39 completely spatially random (CSR) simulations [Baddeley et al., 2015], and included a ‘Flat’ model that estimates SICs independently for each image without hierarchical pooling. Figure 4 illustrates one simulated example; full simulation details are provided in the Supplement.

**Figure 4:**
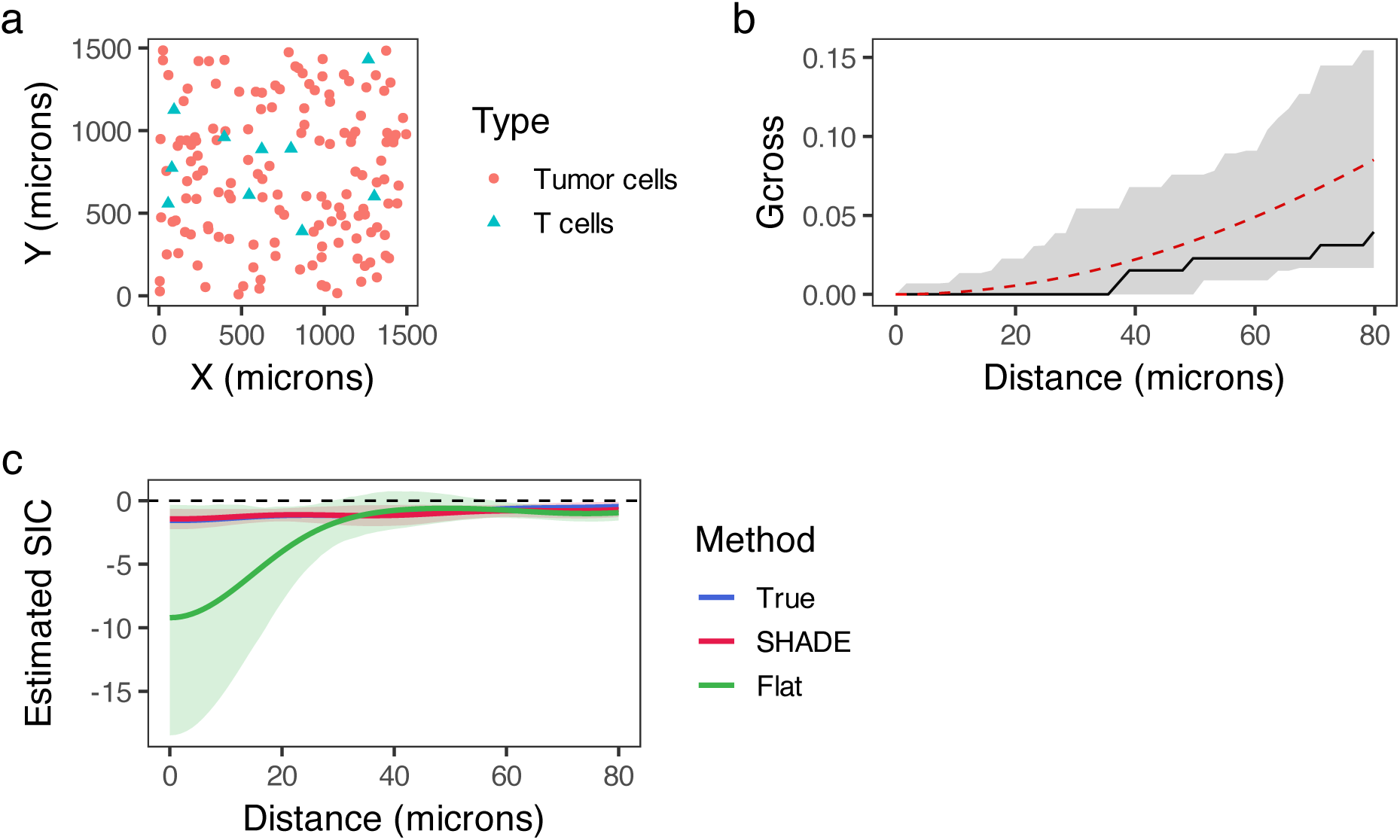
Simulation study comparing spatial analysis methods. (a) Simulated pattern from a responder case showing T cells and tumor cells in a 1500 × 1500 *µ*m^2^ region. (b) *G*-cross analysis for (a), with the observed curve (black), CSR expectation (dashed red), and 95% confidence envelope (gray ribbon) around the null, from 39 simulations. (c) SIC estimates from SHADE (red), a flat model (green), and the ground truth (blue), with 95% credible intervals.

Figure 5 shows that SHADE consistently outperforms both the *G*-cross function and the flat model in detecting true spatial associations, particularly under low T cell or tumor cell densities. While all methods perform well when cell densities are high, the performance of *G*-cross deteriorates markedly under low-density conditions. SHADE maintains robust detection rates even with sparse data and few images per patient, demonstrating the advantage of its hierarchical structure for information sharing across images. This is especially evident in the lower-left and lower-right panels, where SHADE achieves substantially higher detection accuracy compared to the other methods (except for in the case when there is only 1 image per patient and low densities for both types).

**Figure 5:**
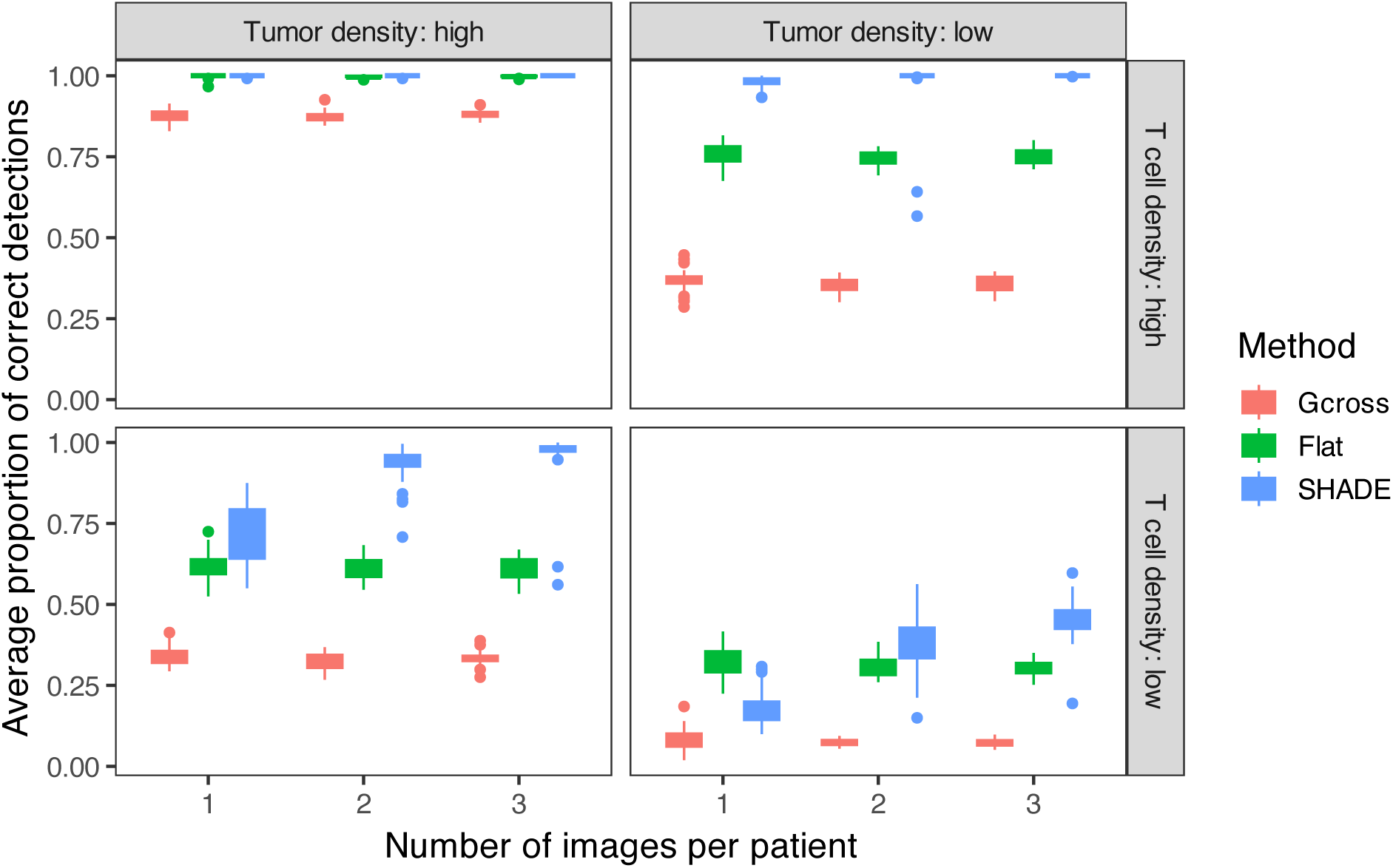
Performance comparison of spatial analysis methods across simulation conditions. Boxplots show the proportion of correctly identified spatial associations at 20, 40, and 60 *µ*m for *G*-cross, Flat, and SHADE models.

## 4 Results: Multiscale inference of directional spatial interactions in colorectal cancer

### 4.1 Description of colorectal cancer dataset

We applied our hierarchical spatial interaction model to a publicly available colorectal cancer (CRC) dataset of multiplexed tumor tissue images from 35 patients [Schürch et al., 2020]. The dataset includes four images per patient, each annotated with single-cell resolution across 16 cell types and 56 protein markers, yielding a multilevel structure of images nested within patients and patients nested within immune phenotype groups: Crohn’s-like reaction (CLR) and diffuse inflammatory infiltration (DII). CLR tumors are typically immune-infiltrated (“hot”), while DII tumors show immune exclusion (“cold”).

For analysis, we focused on the eight most abundant cell types, including reclassification of “stroma” cells as hybrid epithelial-mesenchymal (E/M) cells and “smooth muscle” cells as cancer-associated fibroblasts (CAFs) based on marker expression [Kuburich et al., 2024, Cao et al., 2025]. Target populations were selected for their relevance to anti-tumor immunity (CD8^+^ T cells, memory CD4^+^ T cells, and granulocytes), while source populations included vasculature, tumor cells, CAFs, TAMs, and hybrid E/M cells, reflecting key players in tissue architecture and immune regulation.

For each target cell type, we jointly estimated spatial interaction curves (SICs) with respect to all source types. Detailed data preparation procedures—including quadrature construction, basis function choices for interaction features, and model fitting settings—are provided in the Supplement.

### 4.2 How are specific cell types spatially organized in the tumor microenvironment?

A core question in tumor microenvironment analysis is whether certain cell types tend to cluster near or avoid others, and how these patterns vary with spatial scale. Such spatial relationships reflect underlying mechanisms of attraction, repulsion, communication, or physical constraint, and may provide insight into processes like immune surveillance, tumor evasion, and niche formation. SHADE’s directional SICs allow detection of cell-type-specific clustering and exclusion behaviors across scales.

We extracted image-, patient-, and cohort-level interaction parameters 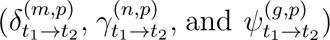 and computed SICs as in Equation 4. Figure 6 shows an example with CTLs as the target cell type and CAFs as a source, highlighting differential organization between patient groups. At ∼75 *µ*m, CTLs in CLR patients tend to cluster around CAFs (log-intensity *>* 0), whereas in DII patients, CTLs appear depleted near CAFs. Patient-level curves reveal substantial within-group heterogeneity.

**Figure 6:**
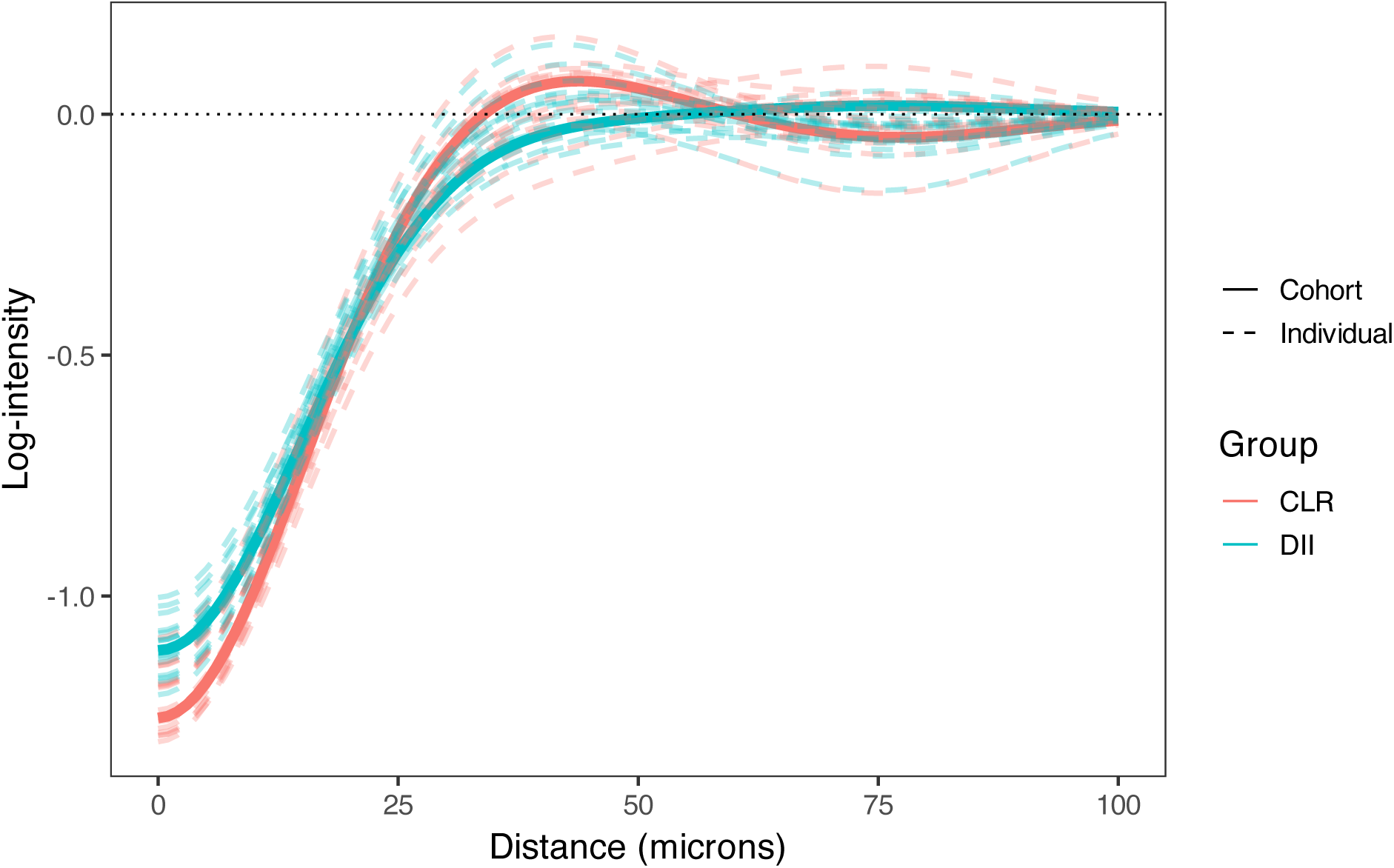
SIC showing the directional association of CAFs (source) with CTLs (target), stratified by patient group. Solid lines show cohort-level estimates for CLR and DII; dashed lines show patient-level SICs. CTLs in CLR patients exhibit midrange clustering around CAFs, while DII patients show neutral to avoidant patterns, indicating group-specific spatial organization.

Across all cell-type pairs, we observed consistent short-range (*<*25 *µ*m) negative associations, likely reflecting physical crowding (see figure in Supplement). At intermediate ranges (25–75 *µ*m), many pairs showed a distinct positive peak, suggestive of clustering, often differing in magnitude between patient groups. Associations at longer distances (*>*75 *µ*m) were generally weaker or absent, indicating limited spatial coordination at larger scales.

### 4.3 How do spatial interaction patterns vary across patients and tissue sections?

Spatial interactions can vary across tumors and even between sections from the same patient, reflecting biologically meaningful heterogeneity in tumor architecture or immune organization. SHADE’s hierarchical structure enables SIC estimation at the image, patient, and cohort levels, allowing us to assess both intra- and inter-patient variability and distinguish conserved from patient-specific spatial patterns.

To assess variability in spatial interaction structure across samples, we summarize between-patient and between-image heterogeneity using median absolute deviation (MAD)-based estimates. These measures are defined in the Supplement. Figure 7 shows heatmaps of the resulting variability estimates for each source–target cell type pair.

**Figure 7:**
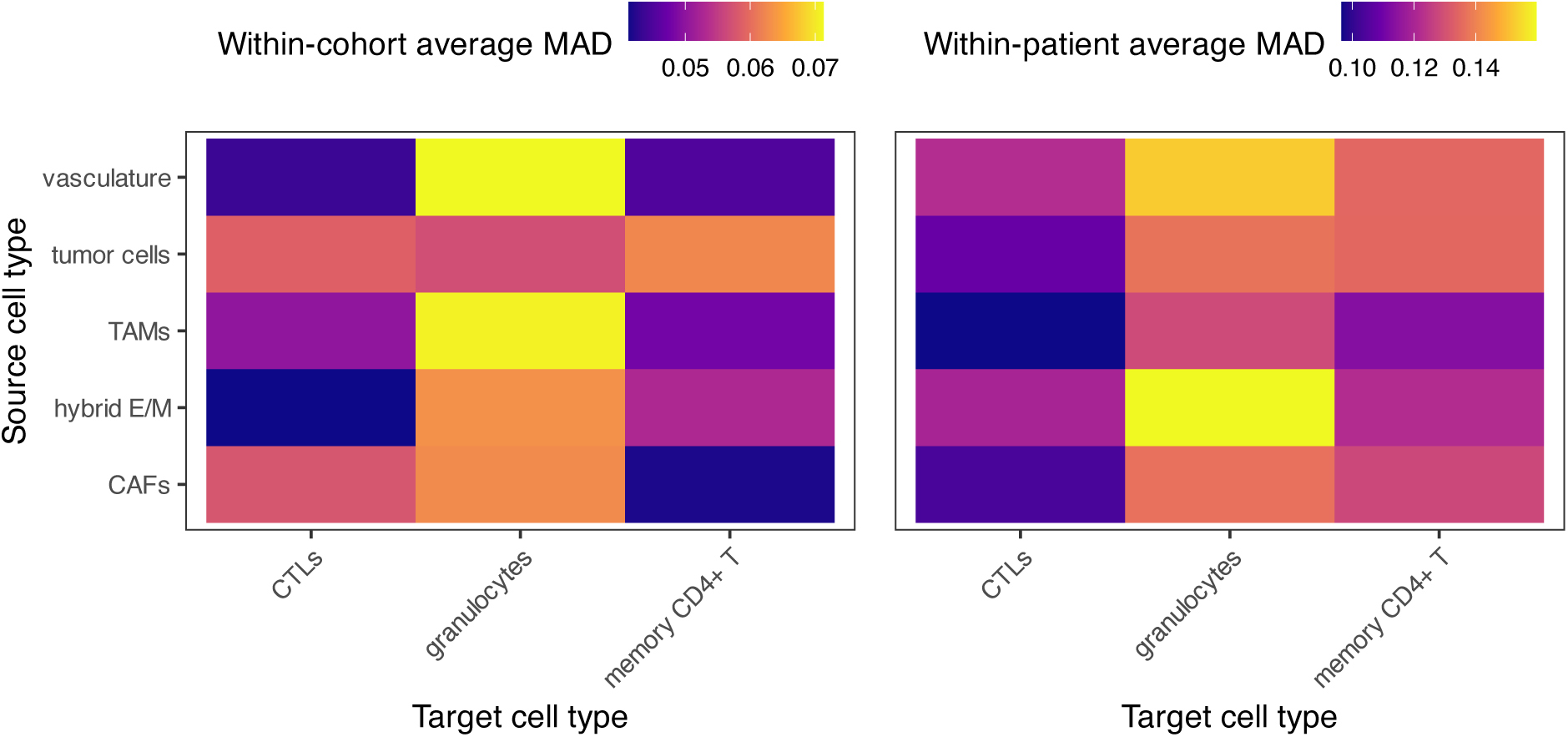
Heatmaps show the median absolute deviation (MAD) of SICs across spatial distances, summarizing two sources of variability: between patients (within cohort) and between images (within patient). Higher values indicate greater heterogeneity in spatial structure.

We observed substantial variability in spatial interaction structure across both patients and images (Figure 7). Notably, granulocytes clustering around TAMs and vasculature exhibited high between-patient heterogeneity. This suggests that granulocyte recruitment and localization near TAMs and vessels may be modulated by patient-specific microenvironmental states, such as differences in angiogenic signaling or myeloid activation. Within patients, granulocyte interactions also showed high heterogeneity, but principally in association with vasculature and hybrid E/M cells. These image-level differences may reflect localized fluctuations in endothelial activation or epithelial plasticity, which may differentially attract or retain granulocytes within specific tissue regions. Examples of patient-level curves for two patients, along with their image-level curves, are shown in Figure 8.

**Figure 8:**
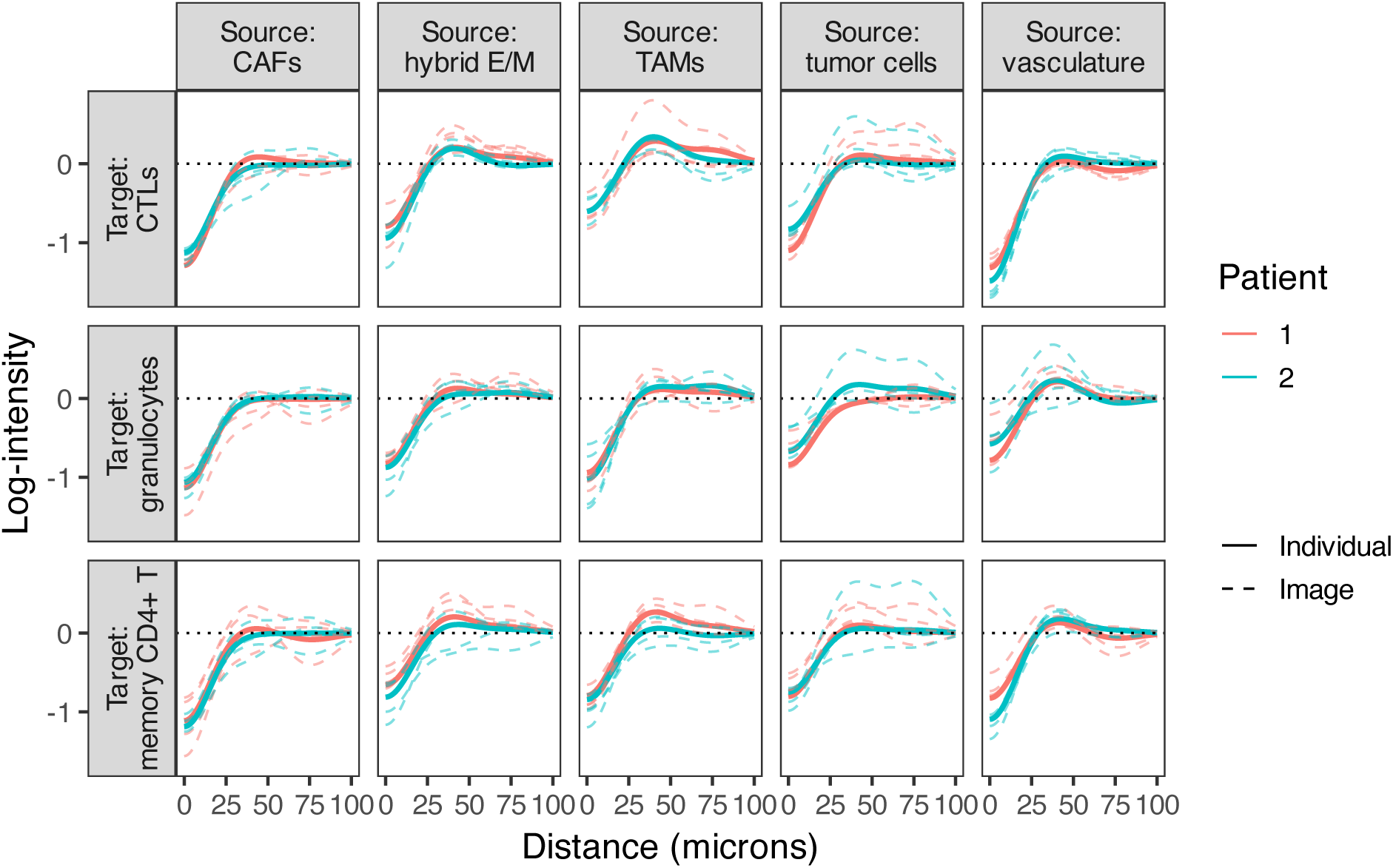
Examples of patient- and image-level SICs for cell type pairs with high within-patient MAD values. Each dotted line represents a SIC estimated from a single tissue section, while each solid line indicates a patient-level SIC.

### 4.4 How do immune and stromal spatial interactions differ between hot and cold tumors?

Distinct tumor immune phenotypes—such as immune-infiltrated versus immune-excluded tumors—are associated with differential immune activity and prognosis. These differences are often accompanied by changes in how immune and stromal cells are spatially arranged in relation to tumor cells and each other. We used SHADE to compare SICs across these patient groups and identify interactions that are enriched or depleted in each context. This allowed us to link differences in spatial structure to known functional differences in immune activity.

We observed several notable differences in SICs between CLR and DII patients (Figure 9). Below, we highlight a subset of interactions where the between-group difference in log-intensity exceeds 0.05 (dashed lines in Figure 9), focusing on plausible underlying biological mechanisms. First, CLR patients showed increased clustering of CTLs and memory CD4+ T cells around vasculature at short spatial ranges, suggesting enhanced immune surveillance and trafficking in these patients. The vasculature serves as a conduit for lymphocyte infiltration into the tumor microenvironment, and vascular normalization in immunologically active tumors has been linked to improved T cell recruitment and anti-tumor immunity [Lanitis et al., 2015, Hendry et al., 2016]. This may reflect a more organized immune response in CLR patients, consistent with better clinical outcomes. In contrast, DII patients exhibited greater clustering of CTLs around tumor cells at all spatial ranges. While this might suggest immune recognition, it may alternatively indicate ineffective or exhausted T cell responses that fail to clear tumor cells. Persistent CTL-tumor colocalization without effective cytolysis has been associated with immune dysfunction and tumor immune escape [Dolina et al., 2021, Raskov et al., 2021].

**Figure 9:**
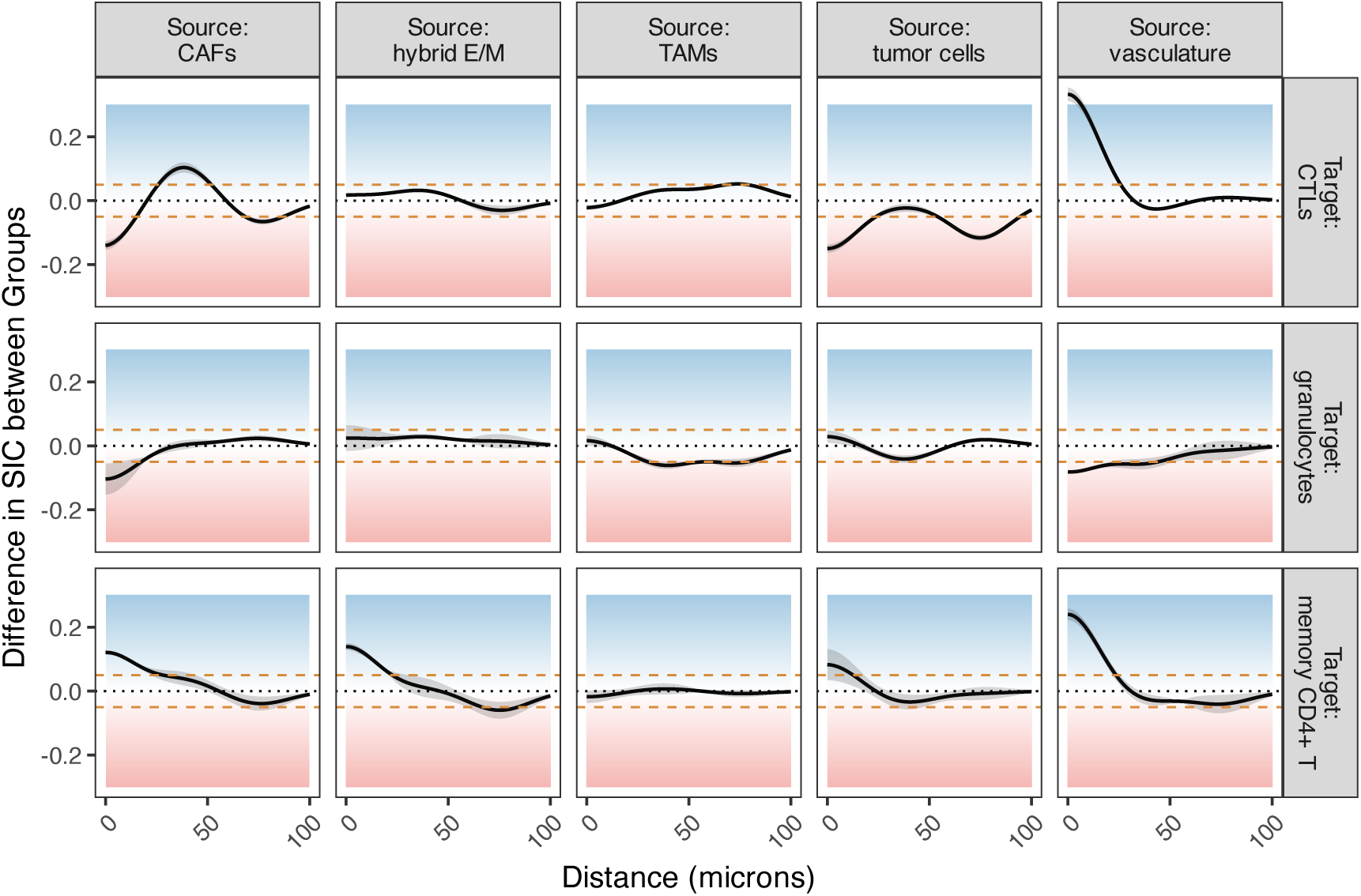
The difference in SICs between the two patient groups. Curves above 0 (in the blue area) indicate that the log-intensity is greater in CLR patients than DII patients; curves in the red area show the reverse.

We also observed distinct patterns in CTL clustering around CAFs: DII patients displayed increased clustering at short and long spatial ranges, whereas CLR patients showed elevated clustering at medium ranges. CAFs are known to mediate immune suppression via direct inhibition of CTLs and by establishing physical and cytokine-mediated barriers [Jenkins et al., 2022, Freeman and Mielgo, 2020]. The long-range clustering in DII patients may reflect exclusionary barriers or CAF-mediated trapping of CTLs, whereas the medium-range interactions in CLR patients could represent transient engagement or immune cell penetration into CAF-dense zones under less suppressive conditions. Lastly, CLR patients demonstrated increased clustering of memory CD4+ T cells at short distances from both CAFs and hybrid E/M cells. This may suggest spatial niches where helper T cells coordinate localized immune responses, potentially promoting anti-tumor immunity or influencing the differentiation state of tumor cells. CD4+ T cells have been implicated in the modulation of EMT processes, and their proximity to hybrid E/M cells may reflect attempts to restrain phenotypic plasticity and metastatic potential [Xie et al., 2025, Milosevic and Östman, 2024].

### 4.5 How well can spatial interactions predict the organization of individual cell types across tumor regions?

Beyond estimating SICs, SHADE predicts the spatial distribution of each target cell type conditional on the source cells, allowing us to assess how well spatial interactions explain observed patterns. This reveals which cell types are more spatially constrained and how predictability varies across images, patients, and tumor subtypes.

For example, in image 47 B from the dataset, we can produce the following predictions of each target cell type, conditional on the source cell types (Figure 10). We can see that the target cell types are quite well predicted, especially CTLs. Likely due to their relative paucity, granulocytes are not quite as well predicted in this image (regions of localization are not as distinct).

**Figure 10:**
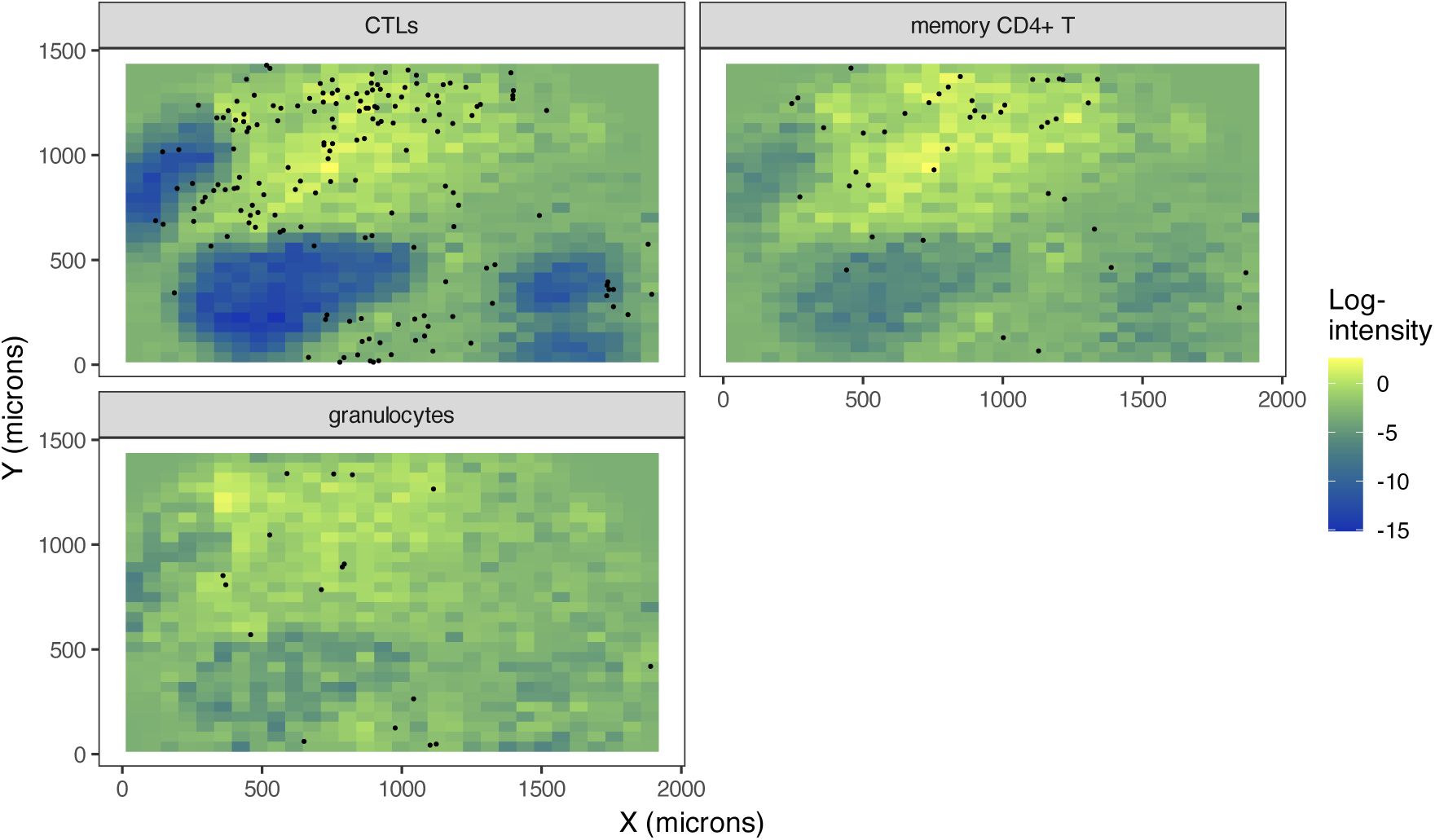
Predicted spatial distributions for target cell types in a representative image (47 B), based on the estimated conditional intensity functions.

We calculated AUCs per cell type and per group (see Supplement). All three target cell types were better predicted in CLR patients than in DII patients, though there was a relatively large amount of variability in image-level AUCs.

### 4.6 Comparing SHADE to traditional ***G***-cross-based spatial clustering analysis

To contextualize our SHADE results, we performed an alternative analysis using the *G*-cross function [Baddeley et al., 2015], a nonparametric estimator of cross-type clustering commonly used in spatial point pattern analysis. This comparison highlights how SHADE differs from traditional methods in its modeling framework and interpretability of results.

For each tissue section, we computed *G*-cross estimates at distances from 0 to 80 microns and extracted values at key distances (20, 40, 60 microns) for comparison. To quantify whether differences in clustering could predict patient group (CLR vs DII), we fitted logistic regression models with *G*-cross as the predictor and used FDR-adjusted p-values to identify significant associations. The resulting estimates are visualized as a heatmap of log-odds ratios (log-ORs), indicating the strength and direction of group differences in clustering at each distance.

The *G*-cross-based analysis (Figure 11) reveals several patterns of cell–cell clustering between CLR and DII patients, but notable differences emerge when we compare these results with SHADE-derived SICs. For instance, TAM–granulocyte interactions showed strong, statistically significant differences at all examined distances (20–60 microns), with higher clustering in DII tumors. This suggests a prominent granulocyte-TAM association in DII tumors according to *G*-cross. However, SHADE did not highlight this pair as a major differential interaction between groups, indicating that the clustering of granulocytes around TAMs may not translate into strong directional interactions when adjusting for other cell types or when modeled hierarchically. Conversely, SHADE identified vasculature as a key source driving differences in clustering of both CTLs and memory CD4+ T cells between CLR and DII groups, reflecting enhanced immune surveillance or trafficking around blood vessels in CLR tumors. This vasculature effect is less evident in the *G*-cross analysis, where vasculature-related interactions showed limited significance and mostly small effect sizes. These discrepancies highlight how SHADE’s joint modeling and multilevel structure may capture spatial dependencies missed by simpler summary statistics like *G*-cross, or may reduce confounding by accounting for other cell types and hierarchical data structure. Conversely, *G*-cross may capture strong pairwise clustering that SHADE’s model de-emphasizes in favor of more complex or conditional dependencies. Overall, this comparison underscores the differences in what each method captures, emphasizing SHADE’s ability to reveal nuanced, context-dependent spatial interactions.

**Figure 11:**
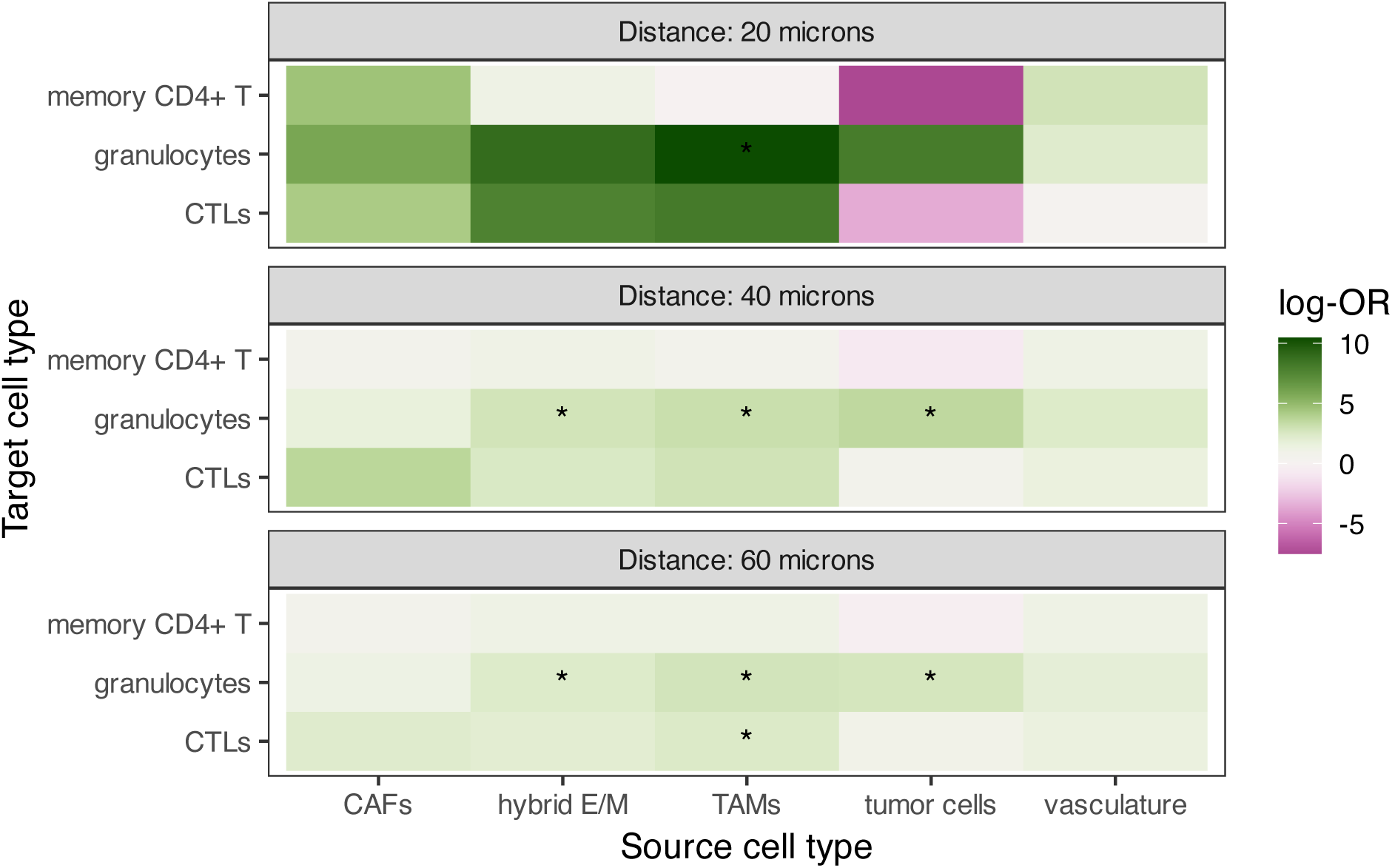
Heatmap of log-odds ratios (log-ORs) from logistic regression models comparing *G*-cross clustering metrics between CLR and DII patient groups, stratified by distance (20, 40, 60 microns). Tiles represent the estimated log-OR for each source–target pair at a given distance, with asterisks indicating significant differences (*p <* 0.05, FDR-adjusted). Green indicates stronger clustering in DII, purple indicates stronger clustering in CLR.

## 5 Discussion

In this work, we introduced SHADE, a Bayesian hierarchical model for quantifying asymmetric spatial interactions in multiplexed imaging data. By modeling the conditional intensity function and estimating spatial interaction curves, our approach enables the identification of directional associations between cell types while accounting for multilevel variation across images, patients, and cohorts.

Through simulation studies, we demonstrated that SHADE effectively captures asymmetric spatial associations while leveraging hierarchical structure to improve estimation. Including multilevel priors improves inference accuracy, especially for short-range interactions, by reducing bias and variance in estimated coefficients. SHADE also demonstrated superior robustness to low cell densities and limited sampling, consistently outperforming *G*-cross and flat models in detecting true spatial interactions across diverse simulation conditions.

SHADE revealed distinct and biologically meaningful patterns of cellular organization in colorectal tumors, highlighting how immune and stromal interactions differ between immune-infiltrated (CLR) and immune-excluded (DII) phenotypes. By estimating directional spatial interaction curves across multiple spatial scales and model hierarchies, we identified key differences in the local microenvironment that may underlie divergent immune states. Notably, CLR tumors exhibited increased short-range clustering of cytotoxic and memory T cells around vasculature, consistent with enhanced immune infiltration via normalized vessels and the formation of perivascular immune niches. In contrast, CTLs in DII tumors showed elevated spatial association with tumor cells across all distances—a pattern that may reflect either failed immune surveillance or dysfunctional retention of exhausted T cells.

Our analysis also uncovered cell-type-specific interaction patterns with stromal components. CTLs in DII patients showed greater clustering near CAFs at both short and long spatial ranges, whereas in CLR patients, CTL–CAF interactions peaked at intermediate distances. Given CAFs’ established role in immune suppression and physical exclusion, these patterns may reflect differential stromal constraint across tumor types. Similarly, CLR patients exhibited increased short-range clustering of memory CD4+ T cells around both CAFs and hybrid E/M cells. These interactions suggest a potentially immunoregulatory or anti-tumor role for helper T cells in CLR tumors, possibly including modulation of EMT plasticity or the stabilization of immune-supportive niches.

Beyond cohort-level differences, we also found substantial heterogeneity in spatial organization across patients and across tissue sections from the same patient. This variability was particularly pronounced in granulocyte interactions with TAMs, vasculature, and hybrid E/M cells, suggesting patient-specific or localized microenvironmental factors influencing spatial localization. Finally, SHADE’s ability to predict target cell distributions from spatial covariates varied by cell type and tumor subtype: immune cell organization was more predictable in CLR tumors, indicating more structured immune environments compared to the more disorganized spatial context of DII tumors. Collectively, these results emphasize the need for multiscale spatial modeling to capture both consistent interaction motifs and the biologically meaningful variability that characterizes tumor immune landscapes.

Additionally, we compared SHADE’s results with *G*-cross function estimates, a standard spatial summary statistic. *G*-cross estimates identified strong clustering of granulocytes around TAMs and hybrid E/M cells, which SHADE did not highlight. Conversely, SHADE revealed significant vasculature-driven clustering of CTLs and memory CD4+ T cells in CLR tumors, a pattern less evident in *G*-cross estimates. This highlights how SHADE’s hierarchical, conditional modeling framework can capture context-dependent interactions that simpler pairwise clustering methods might miss or misrepresent.

A key strength of SHADE is its flexibility in modeling directional interactions across spatial scales, avoiding the restrictive symmetry assumptions of traditional spatial models. Moreover, the use of logistic regression for conditional intensity estimation allows for scalable and stable inference, sidestepping common numerical pitfalls of direct Poisson modeling.

Nonetheless, SHADE has limitations. First, while SICs capture directional spatial association, they remain correlational and cannot determine causality or infer mechanisms of interaction. Second, the SICs reflect spatial dependence rather than molecular signaling pathways, which may limit biological interpretability in some settings. Third, although SHADE supports biologically motivated conditioning structures, exploratory analyses may benefit from modeling both directions of association when directionality is uncertain.

Future extensions of SHADE could incorporate functional covariates—such as marker intensity, proliferation, or exhaustion scores—into the SIC framework, enabling joint analysis of spatial structure and functional state. This would open the door to modeling not just where cells are, but how they behave spatially in context.

Overall, our results highlight the importance of capturing asymmetric and hierarchical structure in spatial models, and position SHADE as a powerful tool for dissecting the complex architecture of tissue microenvironments in cancer and beyond.

