## Supplement for "SHADE: A Multilevel Bayesian Approach to Modeling Directional Spatial Associations in Tissues"

#### 1 Interpreting Example Spatial Interaction Curves

Figure S1 shows two example SICs representative of those estimated from real data. Type 2 cells (cyan) exhibit a strong positive association at short distances, consistent with clustering behavior observed in immune infiltration around tumor cells. In contrast, Type 1 cells (red) show negative association at close range, suggestive of exclusion zones, as might be induced by physical barriers or competitive interactions in the tissue microenvironment.

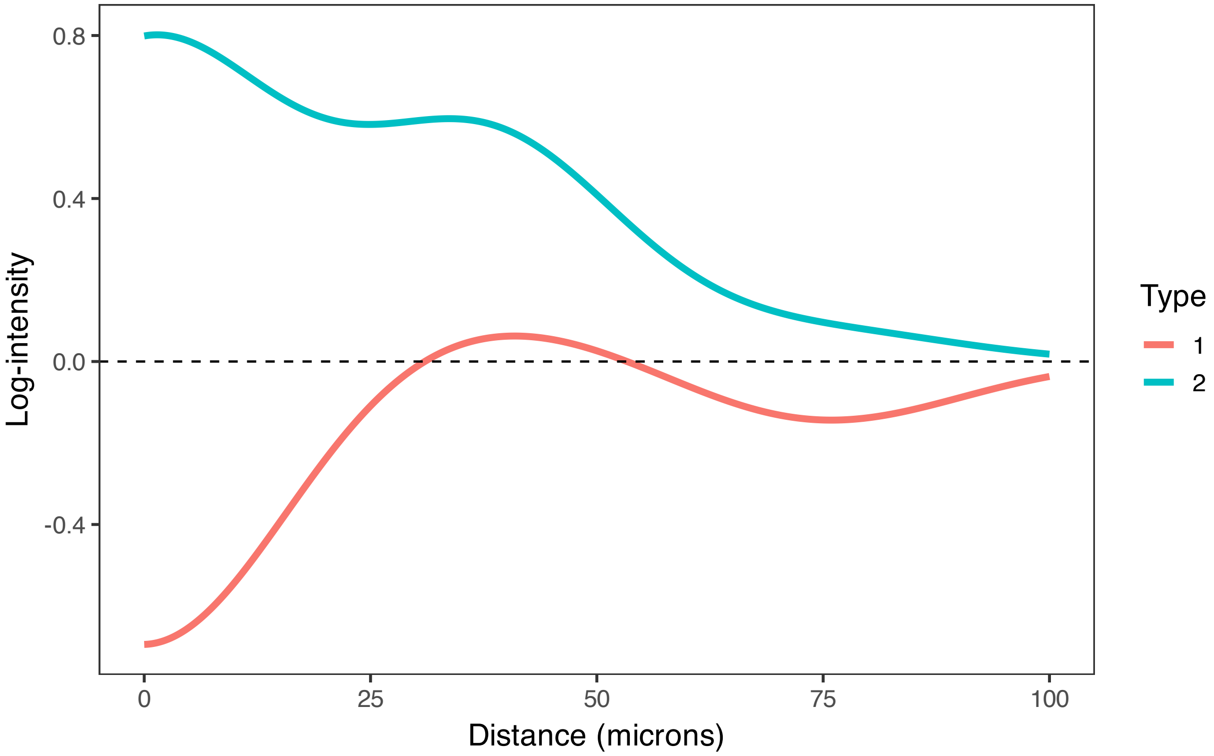


Supplementary Figure S1. Estimated SICs for two source cell types. Type 2 shows a strong positive association with the target population at short distances, while Type 1 exhibits negative association at short range. These contrasting patterns highlight the flexibility of the SIC framework for capturing biologically meaningful spatial structure.

Figure S2 demonstrates how these spatial interactions manifest in tissue space. Panels 1 and 2 show the spatial predictors $\boldsymbol{q}_{A_1}(v)$ and $\boldsymbol{q}_{A_2}(v)$ generated by the individual SICs from Figure S1—each reflecting the localized contribution to the log-intensity of type $B$ due to each source. Each panel is analogous to a summand from the sum on the right hand side of (2). Panel 3 shows the additive combination of these effects, illustrating how multiple source cell types jointly shape a heterogeneous spatial intensity landscape. The target cells are observed to cluster in high-intensity regions and avoid areas of low expected density.

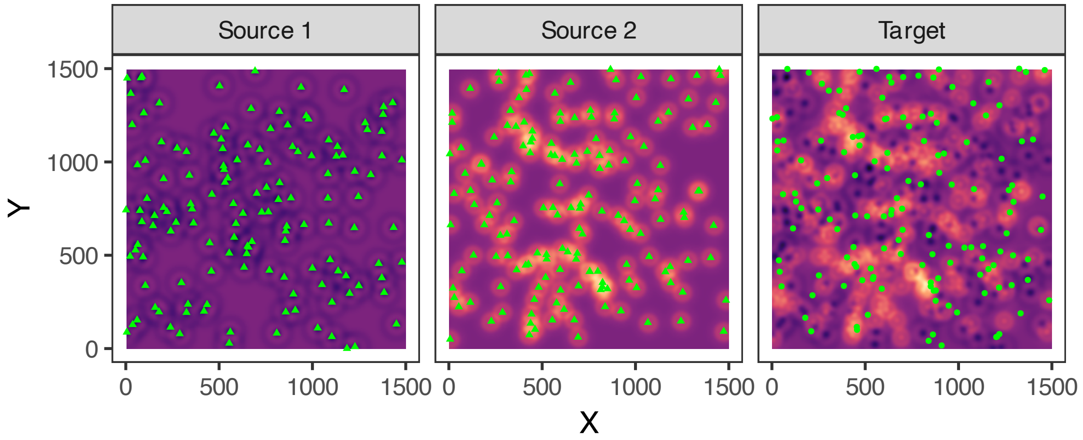


Supplementary Figure S2. Spatial interaction fields implied by SICs from two source cell types (Panels 1 and 2) and their additive combination (Panel 3). The combined field governs the spatial distribution of target cells, which are more likely to occur in regions of elevated log-intensity.

### 2 Extended Methods and Results for Simulation Studies

#### 2.1 Extended Methods for Hyperparameter Studies and Study of Explicit Modeling of Multilevel Structure

##### 2.1.1 Generating Target Points

Target cell locations are drawn from an inhomogeneous Poisson process with intensity:

$$\lambda(v) = \exp\left(\sum_{k=1}^{K}\sum_{p=1}^{P} \boldsymbol{q}_{A_k}^{\top}(v)\boldsymbol{\delta}_{A_k}^{(m)} + \beta_0^{(m)}\right), \tag{1}$$

where $\beta_0^{(m)}$ is a normalization offset to ensure that the expected number of target points matches $N_{\text{points}}$ in image $m$. It is computed as:

$$\beta_0^{(m)} = \log\left(\frac{N_{\text{points}}}{\int_W \exp\left(\sum_{k=1}^{K}\sum_{p=1}^{P} \boldsymbol{q}_{A_k}^\top(v)\boldsymbol{\delta}_{A_k}^{(m)}\right) dv}\right). \tag{2}$$

###### 2.1.2 Default Parameter Settings

Unless otherwise specified, the following defaults were used in simulation experiments:

- **Spatial domain:** $S = 1500$.
- **Number of source cell types:** $K = 2$.
- **Basis functions:** $P = 3$ radial basis functions with bandwidth 15 and support up to 75 microns.
- **Points per cell type per image:** $N_{\text{points}} = 150$.
- **Variance parameters:**
  - $\sigma_{\text{cohort},p} = 0.5$
  - $\sigma_{\text{patient},p} = 0.1$
  - $\sigma_{\text{image},p} = 0.1$
- **Number of patients and images:** variable across experiments.

##### 2.2 Extended Methods for Comparison of Spatial Pattern Detection Accuracy Across Methods and Conditions

###### 2.2.1 Hierarchical Data Generation

We simulate spatial interaction coefficients $\delta^{(m,p)}$ at three hierarchical levels (for image $m$ and basis function $p \in \{1, 2, 3\}$):

$$\psi^{(g,p)} = \begin{cases} -1.5, \ -1.0, \ -0.5 & \text{if group } g = \text{Responder} \\ \ \ 1.5, \quad 1.0, \quad 0.5 & \text{if group } g = \text{Non-responder} \end{cases}$$

$$\gamma^{(n,p)} \sim \mathcal{N}(\psi^{(g(n),p)}, \ \sigma^2_{\text{patient}}), \quad \sigma_{\text{patient}} = 0.1$$

$$\delta^{(m,p)} \sim \mathcal{N}(\gamma^{(n(m),p)}, \ \sigma^2_{\text{image}}), \quad \sigma_{\text{image}} = 0.1$$

Here, $g(n)$ denotes the group of patient $n$, and $n(m)$ denotes the patient corresponding to image $m$.

##### 2.2.2 Spatial Pattern Generation

Cell patterns were generated using spatially varying intensity functions within $1500 \times 1500$ $\mu$m$^2$ observation windows. T cells and B cells were distributed as independent Poisson processes with densities $\lambda_T$ and $\lambda_B$. Tumor cell locations were generated using a spatially varying intensity surface based on T cell locations:

$$\lambda_{\text{tumor}}(s; n) = \exp\left(\beta_0 + \sum_p \delta^{(n,p)} \phi_p(s)\right)$$

where $\phi_p(s)$ represents radial basis functions with $\sigma = 15$ $\mu$m and centered at $\mu = (0, 40, 80)\mu m$. The baseline intensity $\beta_0$ was calibrated to achieve target tumor cell densities, as in (2).

##### 2.2.3 Experimental Design

We employed a factorial design varying the following:

- **Cell Density:** High (150 cells) vs. Low (15 cells) for both T cells and tumor cells.
- **Images per Patient:** 1, 2, or 3 tissue sections.
- **Replication:** 30 independent simulations per condition.

Each simulation included 20 patients per group (40 total), generating datasets with realistic sample sizes for multiplexed imaging studies.

##### 2.2.4 Evaluation Metrics

**SHADE Performance:** We assessed SHADE's ability to recover true spatial interaction curves by constructing pointwise 95% credible bands around posterior estimates. Success was defined as correctly identifying the sign of spatial associations (negative for responders, positive for non-responders) at target distances (20, 40, 60 $\mu$m) using pointwise credible intervals.

**G-cross Comparison:** We implemented envelope tests using 39 simulations of point patterns generated according to complete spatial randomness (CSR) to construct 95% confidence envelopes around the null $G(r)$ function between T cells and tumor cells. Detection success required deviations outside of these null envelopes, with clustering detection for non-responders and repulsion detection for responders.

**Statistical Analysis:** Power was calculated as the average over all images of the proportion of correctly classified interactions at target distances. We compared both methods' sensitivity to varying cell densities and sample sizes.

#### 2.3 Hyperparameter Study 1 - The effect of cell count and dummy point ratio on inference quality

We investigated how inference quality is affected by the number of observed cells per cell type ($N_t$) and the ratio of dummy points to real points ($R_d$) used for quadrature. Simulations were run for all combinations of $N_t \in \{20, 80, 150, 300\}$ and $R_d \in \{0.5, 1, 2, 5, 10\}$, with five replicates per setting. Each dataset contained three cell types, with the third serving as the target for spatial interaction estimation. The number of patient groups was fixed at 1, with 40 patients per group and two images per patient. All other simulation parameters were set to defaults.

Average root mean squared error (RMSE) was computed for spatial interaction coefficients at the image ($\delta^{(m,p)}_{t_1 \to t_2}$), patient ($\gamma^{(n,p)}_{t_1 \to t_2}$), and cohort ($\psi^{(g,p)}_{t_1 \to t_2}$) levels. As shown in Figure S3, RMSE for image-level coefficients decreased consistently with increasing $R_d$, suggesting that finer quadrature grids improve inference at the image level. This trend held across all values of $N_t$.

In contrast, inference quality for patient- and cohort-level coefficients slightly worsened when $R_d$ exceeded 2–5, particularly at very low or very high values of $N_t$. This suggests diminishing returns—and potential instability—for higher-level parameter estimation when dummy point counts are excessively large. The magnitude of this effect was modest but visible (note the log scale on the RMSE axis).

To further investigate these trends, we analyzed RMSE stratified by spatial scale (short, medium, long) at each hierarchical level (Figure S4). Improvements in image-level inference with increasing $R_d$ were most pronounced at longer interaction distances. For higher-level parameters, performance remained relatively stable across spatial scales but showed a slight increase in RMSE at short ranges when dummy point ratios were high.

A representative cohort-level SIC estimate is shown in Figure S5, illustrating how estimation accuracy varies with $N_t$ and $R_d$. Inference was notably poorer at low cell counts, with increased bias and wider credible intervals. Optimal performance—reflected in reduced bias and uncertainty—was observed at moderate $N_t$ (80–300) and a wide range of dummy point ratios, consistent with the RMSE patterns observed in Figure S3.

#### 2.4 Hyperparameter Study 2 - The effect of number of patients and images per patient on inference quality

Next, we examined the sensitivity of inference quality to the total number of patients as well as the number of images per patient. Here, we set the number of points per type to 150 and the ratio of dummy points to actual points to 2, while keeping the rest of the default parameters the same, as outlined in the Supplement. We then simulated 15 realizations each of every combination of the number of images per patient $N_{\text{images per patient}}$

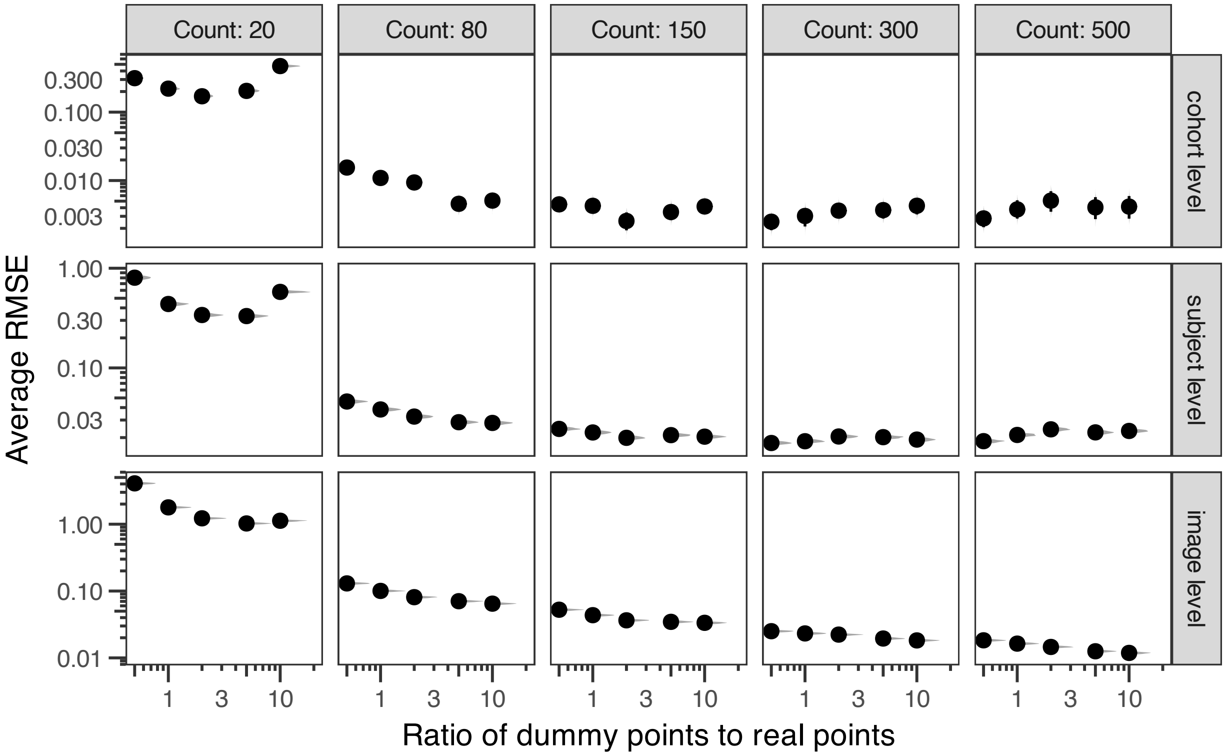


Supplementary Figure S3. Average RMSE for spatial interaction coefficients across hierarchical levels: image-level ($\delta^{(m,p)}_{t_1 \to t_2}$), patient-level ($\gamma^{(n,p)}_{t_1 \to t_2}$), and cohort-level ($\psi^{(g,p)}_{t_1 \to t_2}$), under different numbers of cells per type and dummy-to-real point ratios.

and the total number of patients $N_{\text{patients}}$, where $N_{\text{images per patient}} \in \{1, 2, 4\}$ and $N_{\text{patients}} \in \{10, 20, 40\}$, according to our simulation procedure described in the Supplement.

We found some interesting trends in the average RMSE as the number of patients per patient group increased. For cohorts that only had one image per patient, average RMSE decreased as the number of patients increased (Figure S6). However, for patients that had 2 images, the average RMSE was highest for the highest number of patients, which we found to be a counterintuitive finding. Furthermore, for patients with 4 images, having a cohort with 40 patients was associated with the lowest RMSE, as expected, though the second-lowest was, unexpectedly, having a cohort with 10 patients, rather than 20.

For $\delta^{(m,p)}_{t_1 \to t_2}$ coefficients, longer-range coefficients seemed to have the lowest average RMSE (Figure S7), while close-range coefficients had the worst quality. RMSE, however, was very close to constant across the number of patients, though it did decrease slightly as the number of images increased. RMSE for $\gamma^{(n,p)}_{t_1 \to t_2}$ coefficients decreased as the number of images increased, though, as we noticed earlier, the RMSE in this case has a nonlinear relationship with the number of patients. The RMSE of $\psi^{(g,p)}_{t_1 \to t_2}$ coefficients exhibited the same counterintuitive nonlinear association with number of patients, though in a more pronounced way.

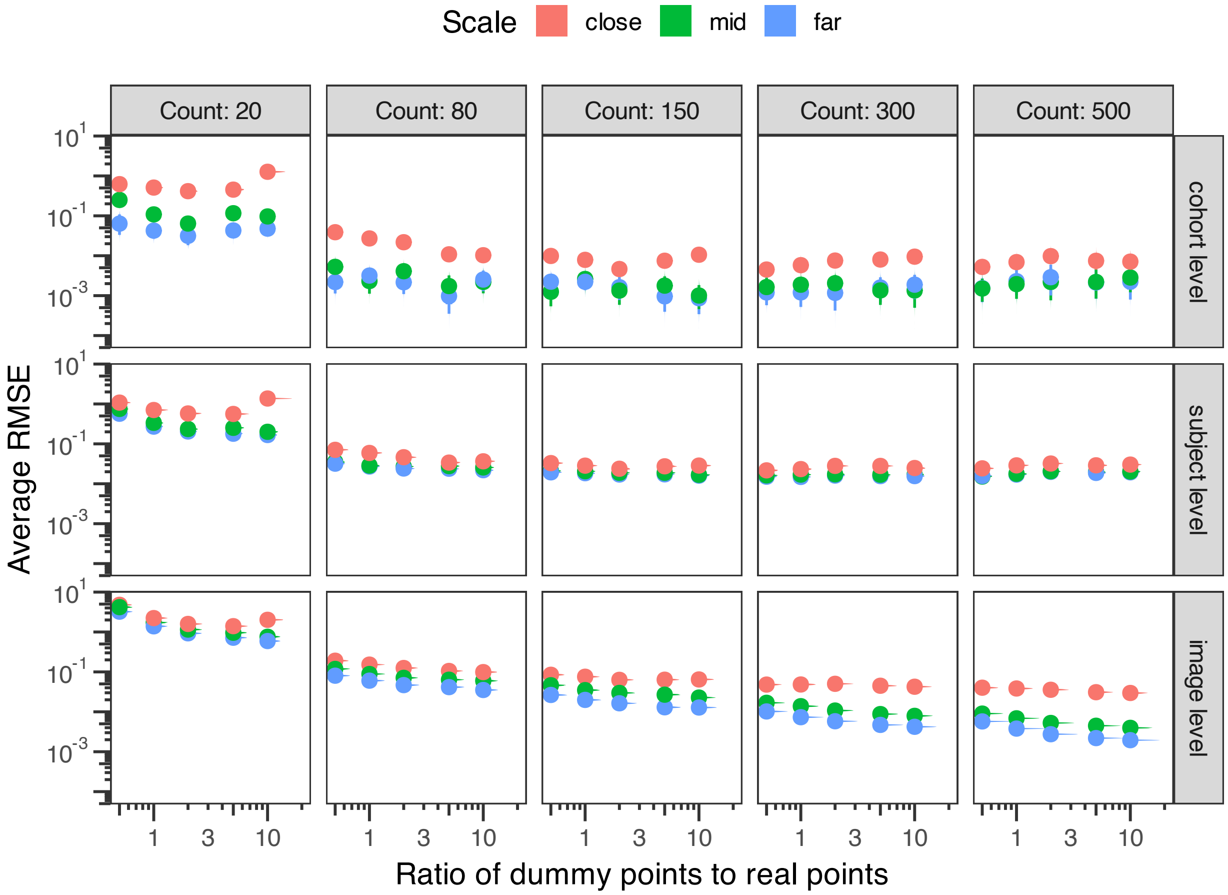


Supplementary Figure S4. RMSE for spatial interaction coefficients across spatial scales, shown separately for each hierarchical level: $\delta^{(m,p)}_{t_1\to t_2}$, $\gamma^{(n,p)}_{t_1\to t_2}$, and $\psi^{(g,p)}_{t_1\to t_2}$. Results are shown for varying cell counts and dummy point ratios.

Finally, we demonstrate an example of an estimated global SIC from this simulation study (Figure S8). Here, we can see that the variance of the SIC estimate decreases both as the number of patients increases and the the number of images increases, which matches our expectations. However, we can also see that there is a slight amount of persistent bias at close range for simulations in which the number of patients equals 40 - this may be indicative of too much shrinkage during estimation and may be the reason for the nonlinear association of RMSE with number of patients.

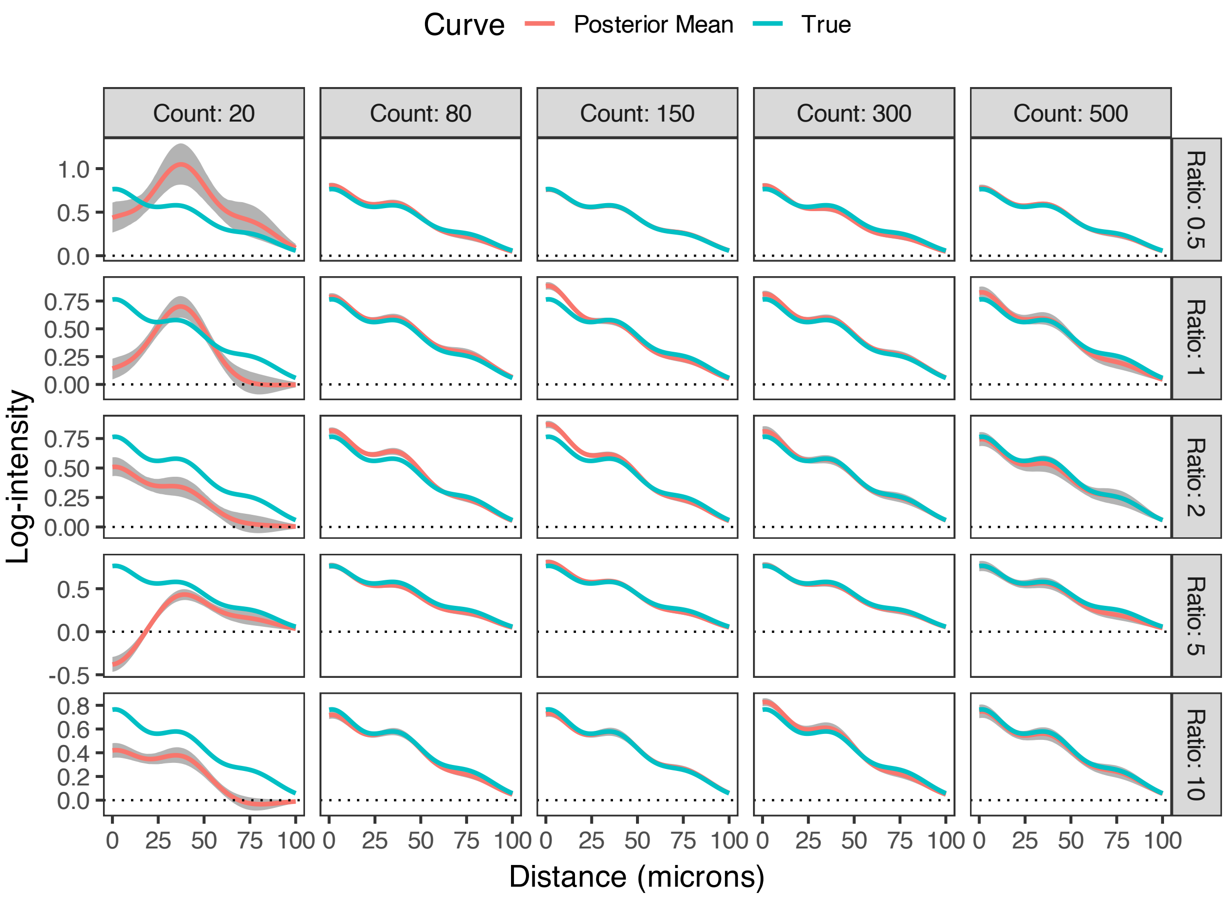


Supplementary Figure S5. Example of estimated cohort-level SIC from dummy point simulations.

### 3 Extended results for colorectal cancer analysis

#### 3.1 Detailed description of colorectal cancer dataset and model preparation

The colorectal cancer (CRC) dataset used in this study is a publicly available collection of multiplexed tumor tissue images from 35 patients (Schürch et al., 2020). Each patient contributed four images, each derived from separate biopsies, yielding a total of 140 images. Images were annotated with single-cell resolution across 16 cell types and 56 protein markers, resulting in a multilevel structure: images nested within patients, patients nested within two immune phenotype groups—Crohn's-like reaction (CLR) and diffuse inflammatory infiltration (DII).

The dataset contains approximately 200,000 cells. For analysis, we focused on the eight most abundant cell types. Cell labels were refined to better reflect marker-based characterization: "stroma" cells were reclassified as hybrid epithelial-mesenchymal (E/M) cells based on co-expression of cytokeratin and vimentin (Kuburich et al., 2024), while "smooth muscle" cells were relabeled as cancer-associated fibroblasts (CAFs)

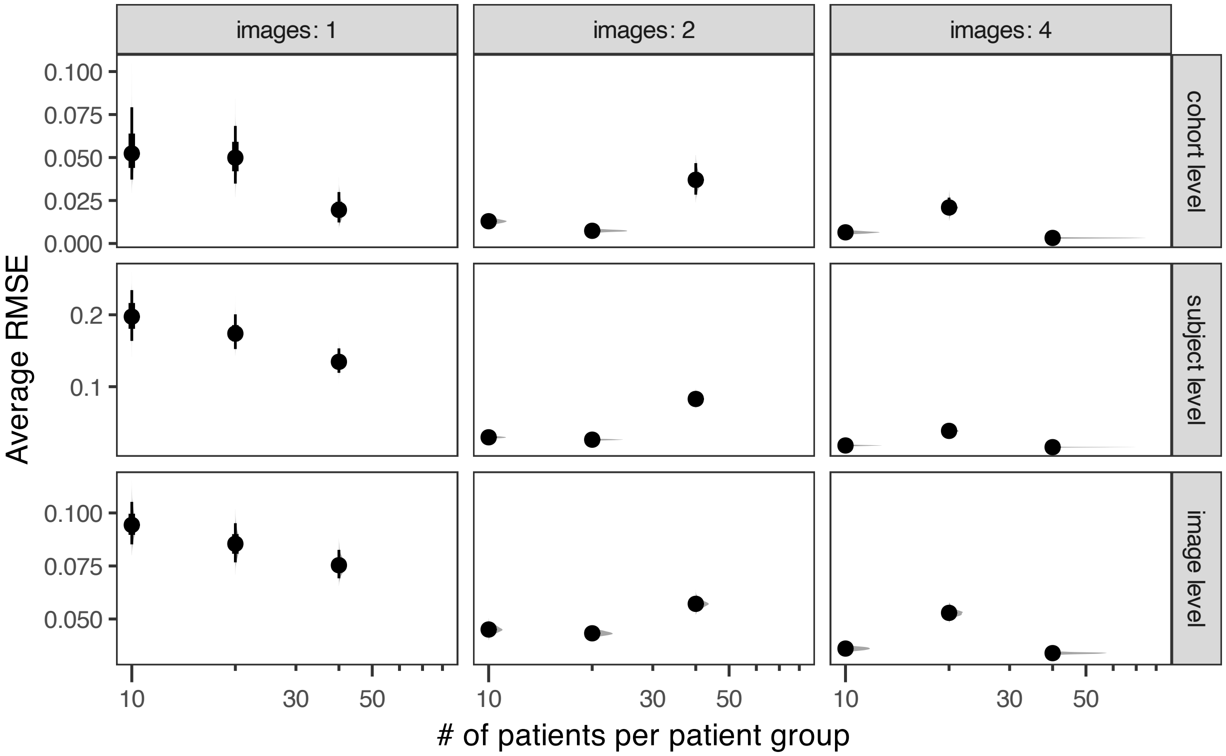


Supplementary Figure S6. Average RMSE for spatial interaction coefficients as a function of dataset size, varying the number of patients and number of images per patient. Results are shown for image-level ($\delta_{t_1 \to t_2}^{(m,p)}$), patient-level ($\gamma_{t_1 \to t_2}^{(n,p)}$), and cohort-level ($\psi_{t_1 \to t_2}^{(g,p)}$) parameters.

due to expression of $\alpha$-SMA and vimentin (Cao et al., 2025). Additional cell types included CD163$^+$ macrophages (TAMs), CD8$^+$ T cells, granulocytes, memory CD4$^+$ T cells, tumor cells, and vasculature.

We selected target populations (CD8$^+$ T cells, memory CD4$^+$ T cells, and granulocytes) based on their functional relevance to anti-tumor immunity, and source populations (vasculature, tumor cells, CAFs, TAMs, hybrid E/M cells) based on their roles in tissue architecture and immune modulation. Vasculature structures infiltration pathways, tumor cells and CAFs contribute to immune exclusion, TAMs modulate local inflammation and immune suppression, and hybrid E/M cells may influence spatial dynamics through motility and stromal interactions.

For each target cell type, we constructed a quadrature scheme by generating 1,000 dummy points per image per cell type. To capture distance-dependent spatial interactions, we constructed interaction features $\boldsymbol{q}_{A_k}(v)$ using a set of three radial basis functions $\phi_p$. Data preprocessing included normalization of coordinates and preparation of covariates and interaction features.

Model fitting was performed using variational inference with 1,000 posterior draws after fitting. SICs were estimated jointly for all source cell types with respect to each target, providing interpretable, distance-

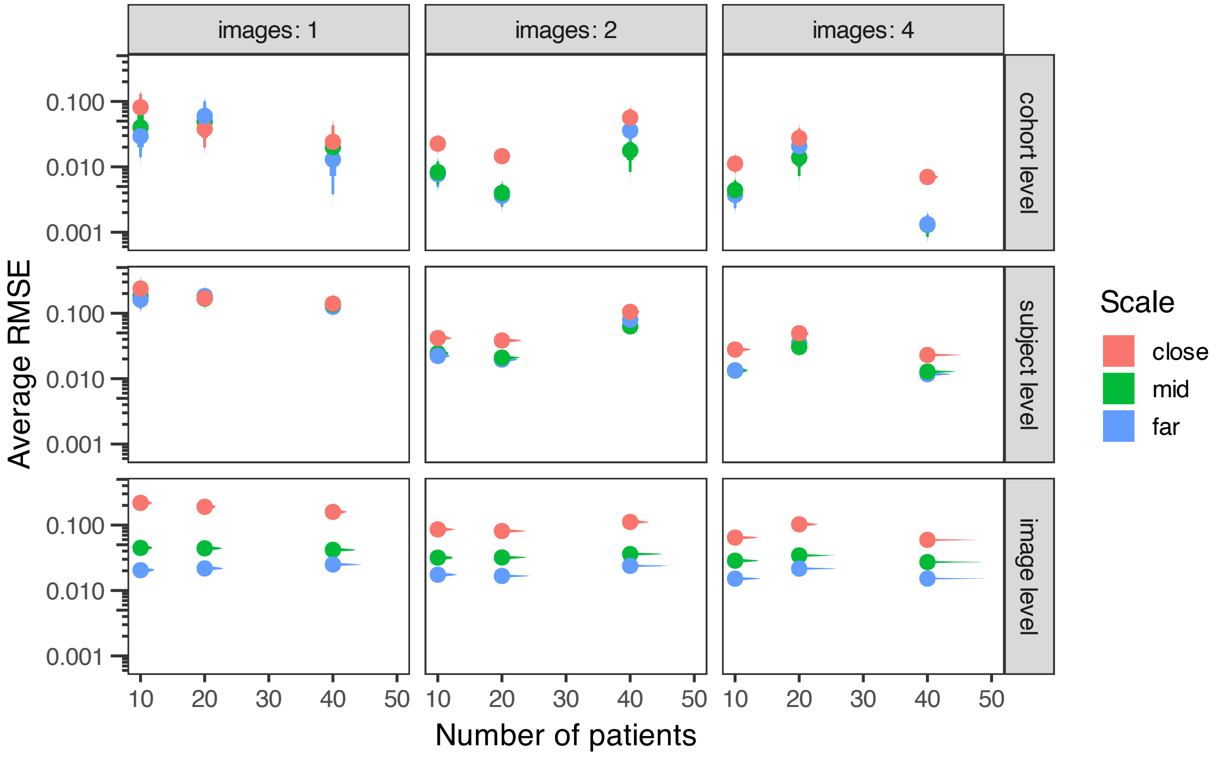


Supplementary Figure S7. Average RMSE across spatial scales for spatial interaction coefficients at the image ($\delta^{(m,p)}_{t_1 \to t_2}$), patient ($\gamma^{(n,p)}_{t_1 \to t_2}$), and cohort ($\psi^{(g,p)}_{t_1 \to t_2}$) levels, across varying numbers of patients and images per patient.

resolved summaries of spatial associations.

#### 3.2 Supplementary Figures

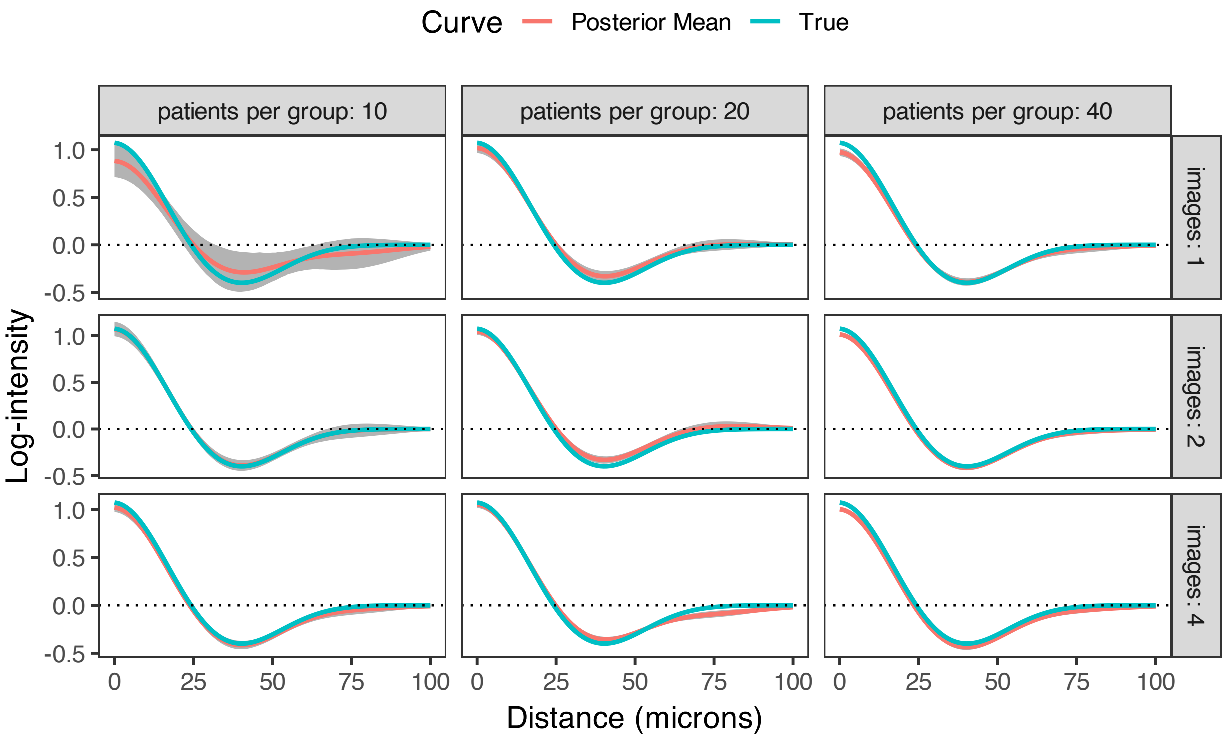


Supplementary Figure S8. Example of an estimated cohort-level SIC from the dataset size simulation study, illustrating how increasing the number of patients and images per patient reduces the variance in SIC estimates.

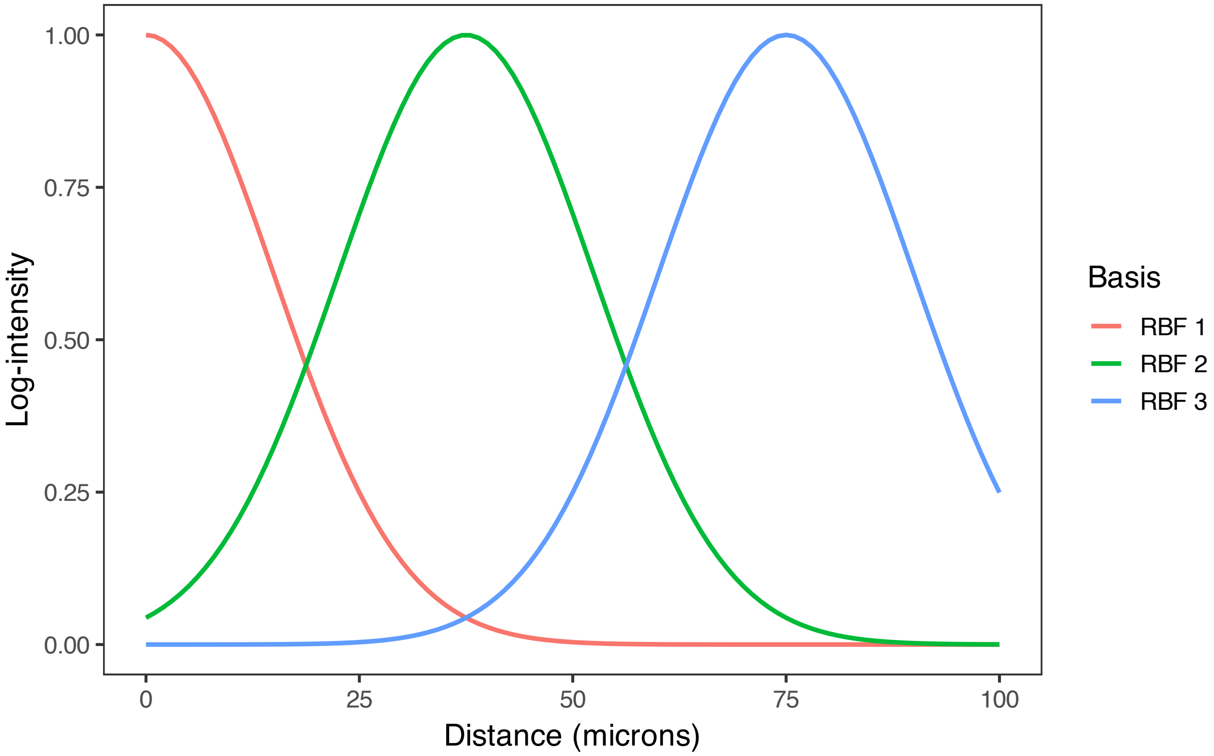


Supplementary Figure S9. Radial basis functions $\phi_p(s)$ used to compute distance-based interaction features $\boldsymbol{q}_{A_k}(v)$ in the CRC analysis. These basis functions define the spatial resolution of the SICs.

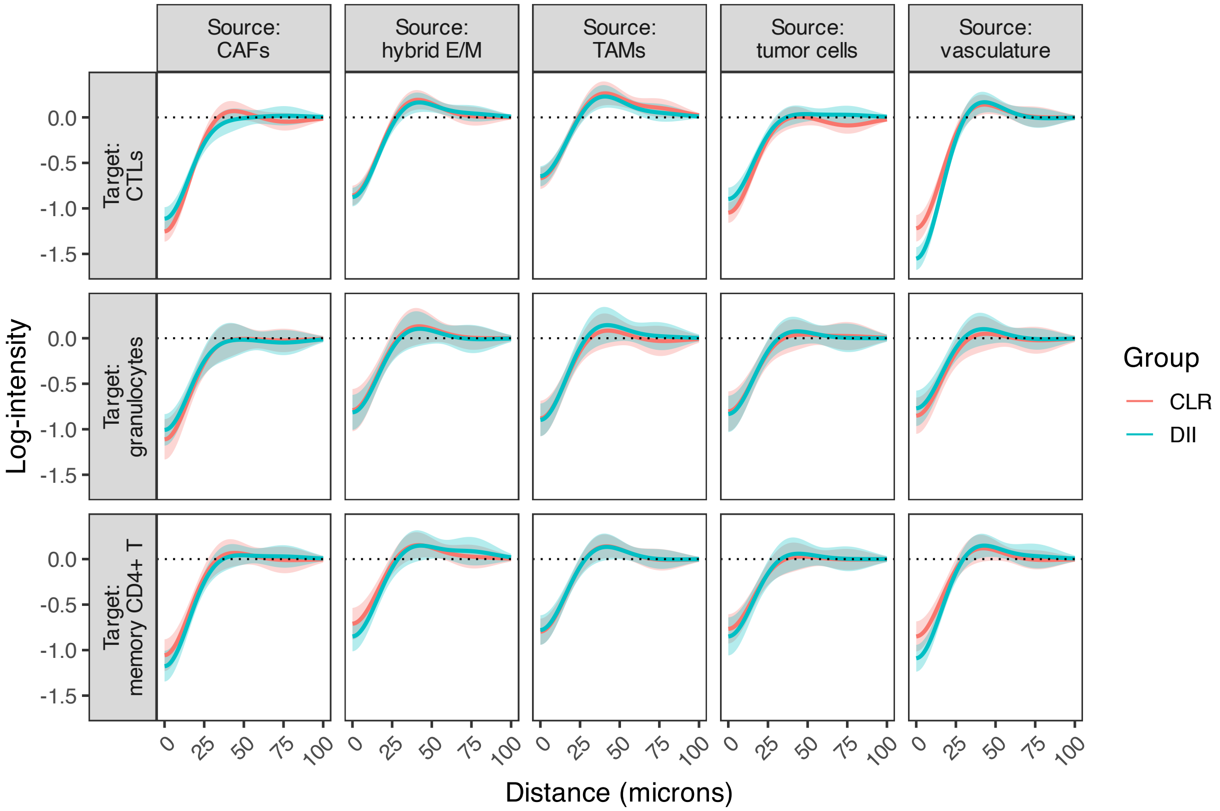


Supplementary Figure S10. Cohort-level SICs ($\psi_{t_1 \to t_2}^{(g,p)}$) estimated for all source–target cell type pairs in the CRC dataset, stratified by CLR and DII patient groups.

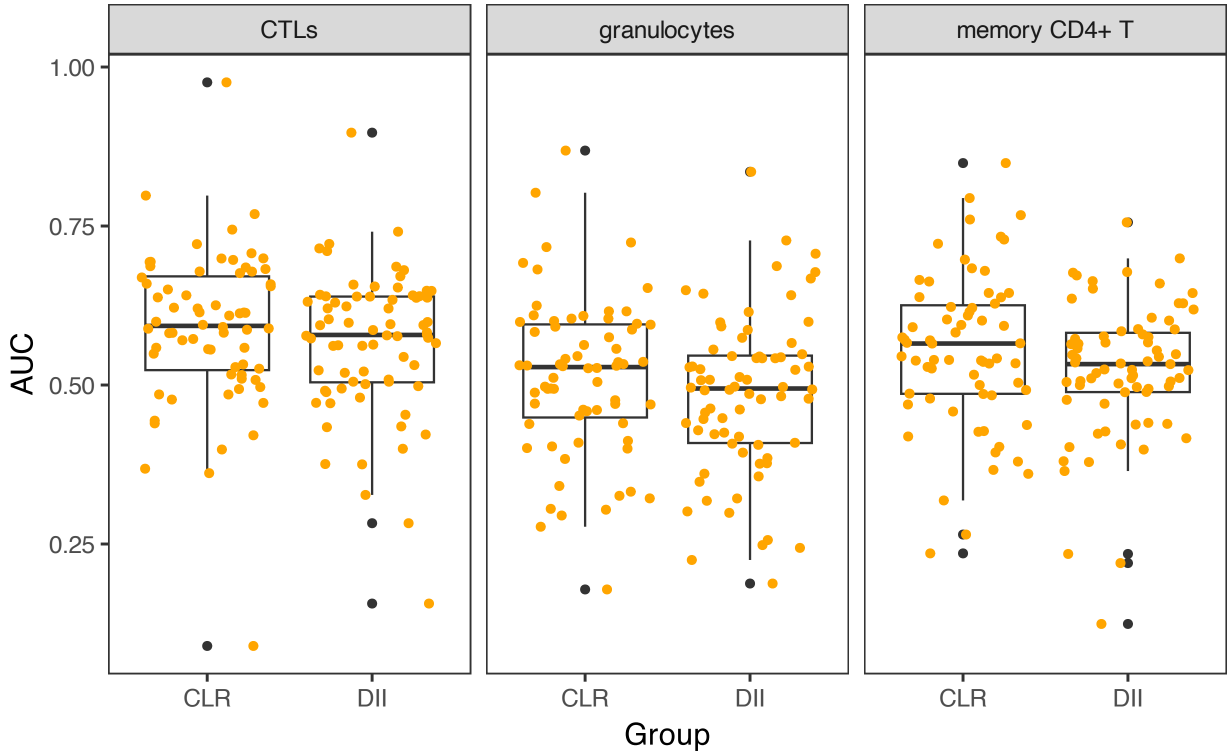


Supplementary Figure S11. Distribution of AUCs by cell type and patient group, evaluating prediction performance of SHADE's conditional intensity model when predicting the spatial organization of each target cell type based on the spatial distribution of all other types.

### 4 Priors for Variance Parameters

We place half-normal priors on the standard deviations of the spatial interaction coefficients at each level of the hierarchy:

$$\sigma_{\text{cohort},p} \sim \text{Half-Normal}(\xi_p, \xi_p), \tag{3}$$

$$\sigma_{\text{patient},p} \sim \text{Half-Normal}(1.5, 10), \tag{4}$$

$$\sigma_{\text{image},p} \sim \text{Half-Normal}(1, 10). \tag{5}$$

where is a user-specified location/scale (in our CRC analysis, we used $\xi_p \in \{5, 3, 1\}$).

### 5 Quantifying Variability in Spatial Interactions

We define SICs hierarchically across cohort, patient, and image levels. For a given source–target cell type pair $(A, B)$, to capture hierarchical variation, we define *patient-level* and *image-level* SICs as:

$$\text{SIC}^{(n)}_{A_k \to B}(s) = \sum_{p=1}^{P} \gamma^{(n,p)}_{A_k} \phi_p(s) \tag{6}$$

$$\text{SIC}^{(m,n(m))}_{A_k \to B}(s) = \sum_{p=1}^{P} \delta^{(m,p)}_{A_k} \phi_p(s) \tag{7}$$

where $\phi_p(s)$ are spline basis functions, and $\gamma^{(n,p)}_{A,B}$ and $\delta^{(m,p)}_{A,B}$ are patient- and image-level coefficients, respectively. To quantify heterogeneity in SICs across patients and images, we compute robust measures of variability at each spatial distance $s$, and summarize them by taking the median across all distances. Specifically:

**Between-patient (within-cohort) variability.** Let $\mathcal{C}$ index cohorts, and let $\mathcal{N}_c$ denote the set of patients belonging to cohort $c \in \mathcal{C}$. For each spatial distance $s$, we compute the cohort-level median SIC as:

$$\overline{\text{SIC}}^{(c)}_{A_k \to B}(s) = \text{median}_{n \in \mathcal{N}_c} \left( \text{SIC}^{(n)}_{A_k \to B}(s) \right)$$

The between-patient (within-cohort) variability for a given cell type pair $(A, B)$ is then defined as:

$$\text{MAD}_{\text{patient}}(A, B) = \text{median}_s \left\{ \text{MAD}_n \left[ \text{SIC}^{(n)}_{A_k \to B}(s) - \overline{\text{SIC}}^{(c(n))}_{A_k \to B}(s) \right] \right\} \tag{8}$$

**Between-image (within-patient) variability.** Similarly, for each patient $n$, we compute the patient-level median SIC:

$$\overline{\mathrm{SIC}}^{(n)}_{A_k \to B}(s) = \mathrm{median}_{m:n(m)=n} \left( \mathrm{SIC}^{(m,n)}_{A_k \to B}(s) \right)$$

The between-image variability is defined as the MAD of image-level SICs from the patient median:

$$\mathrm{MAD}_{\mathrm{image}}(A, B) = \mathrm{median}_s \left\{ \mathrm{MAD}_m \left[ \mathrm{SIC}^{(m,n(m))}_{A_k \to B}(s) - \overline{\mathrm{SIC}}^{(n(m))}_{A_k \to B}(s) \right] \right\} \tag{9}$$

These robust statistics provide interpretable summaries of spatial heterogeneity at each level of the hierarchy while mitigating sensitivity to outliers and small-sample variability. We report these values for each cell type pair and visualize them using heatmaps (Figure 7).
